# Adaptable, Turn-On Monobody (ATOM) Fluorescent Biosensors for Multiplexed Detection in Cells

**DOI:** 10.1101/2023.03.28.534597

**Authors:** Harsimranjit Sekhon, Jeung-Hoi Ha, Maria F. Presti, Spencer B. Procopio, Paige O. Mirsky, Anna M. John, Stewart N. Loh

## Abstract

A grand challenge in biosensor design is to develop a single molecule, fluorescent protein-based platform that can be easily adapted to recognize targets of choice. Conceptually, this can be achieved by fusing a small, antibody-like binding domain to a fluorescent protein in such a way that target binding activates fluorescence. Although this design is simple to envision, its execution is not obvious. Here, we created a family of adaptable, turn-on monobody (ATOM) biosensors consisting of a monobody, circularly permuted at one of two positions, inserted into a fluorescent protein at one of three surface loops. Multiplexed imaging of live human cells co-expressing cyan, yellow, and red ATOM sensors detected the biosensor targets (WDR5, SH2, and hRAS proteins) that were localized to the nucleus, cytoplasm, and plasma membrane, respectively, with high specificity. ER- and mitochondria-localized ATOM sensors also detected ligands that were targeted to those organelles. Fluorescence activation involved ligand-dependent chromophore maturation with fluorescence turn-on ratios of >20-fold in cells and up to 100-fold *in vitro*. The sensing mechanism was validated with three arbitrarily chosen monobodies inserted into jellyfish as well as anemone lineages of fluorescent proteins, suggesting that ATOM sensors with different binding specificities and additional colors can be generated relatively quickly.

## Introduction

Single molecule, fluorescent protein-based biosensors (FPBs) are transforming the study of biochemical processes in cells and organisms. These proteins typically consist of one or more fluorescent proteins (FPs) from jellyfish, coral, anemone, or other sea creature, fused to a receptor protein that confers specificity. Binding of the target analyte to the receptor domain triggers a conformational change in the protein that produces either a ratiometric fluorescence change (via Förster resonance energy transfer) or an increase in fluorescence intensity.

FPBs possess several qualities have made them among the most desirable sensor type in biology. First, their full genetic encoding allows them to be readily introduced into cells and used in many organisms. Second, they require no cofactors or reagents for operation. Lastly, their fluorescence output modes make them useful for applications that demand quantification of ligand concentration as well as high sensitivity. Ratiometric signal establishes a response that is theoretically independent of sensor concentration and therefore consistent from cell to cell, and fluorescence turn-on provides high sensitivity (at the expense of precise quantification).

Perhaps the most influential example of an FBP is the genetically encoded calcium indicator (GECI) family of calcium sensors^1–3^. The output domain of GECIs is an FP that has generally been circularly permuted with its new amino and carboxy termini positioned in a β-strand that packs against the chromophore. The resulting structural defect allows the chromophore to mature but in a protonated state (presumably due to its increased solvent accessibility) that is consequently nonfluorescent or has a decreased fluorescence lifetime^4^. The recognition domain consists of calmodulin (CaM) and one of its cognate binding peptides (e.g., ckkap^5,6^ or RS20^7,8^) fused to each terminus of the permuted FP. On binding Ca^2+^, CaM associates with the peptide, closing around the permutation site and permitting the chromophore to deprotonate. By virtue of their strong and reversible fluorescent turn-on and rapid kinetics, GECIs have revolutionized the study of calcium signaling in cells. This sensing mechanism, however, has proven difficult to translate to recognition domains other than CaM. While successful examples exist^3^, each sensor was the result of a new protein engineering effort whose outcome was far from guaranteed.

A major task facing inventors of FPBs is thus to create a protein switch that can be readily modified to sense new targets as they arise. An intuitive approach for doing so is to fuse an adaptable, antibody-like protein to an FP such that ligand binding activates fluorescence. Several engineered binding scaffolds^9^ are available for this purpose, including nanobodies^10^, DARPins^11,12^, affibodies^13^, de novo designed helical bundles^14,15^, and monobodies (MBs) based on the 10^th^ human fibronectin type III domain^16^. These domains are advantageous because they are small (<15 kDa) and amenable to affinity maturation by *in vitro* and *in vivo* methods. Unlike CaM, however, they do not appreciably change conformation on binding. It therefore remains a formidable challenge to discover mechanisms by which ligand binding to the recognition domain translates to an increase in fluorescence of the FP domain.

Here, we created a family of adaptable turn-on monobody (ATOM) fluorescent sensors that are genetically encoded and can be easily modified to recognize a variety of ligands. We took advantage of a serendipitous observation from our earlier study in which we inserted a circularly permuted FK506 binding protein into the loop connecting strands β10 and β11 of a YFP variant (YFP contains 11 β-strands)^17^. The YFP had itself been circularly permuted between strands β9 and β10. The resulting fusion protein exhibited a 35-fold turn-on of yellow fluorescence upon addition of FK506 or its analog rapamycin. We hypothesized that inserting a circularly permuted MB between strands β10 and β11 of circularly permuted YFP, and possibly between other strands as well, would produce the same turn-on effect on binding the ligand to which the MB was raised. We further surmised that sensors for new targets could be generated by swapping out one MB for another, and that this mechanism might be transferrable to other colors and lineages of the FP.

We report the generation of cyan (c-ATOM), yellow (y-ATOM), and red (r-ATOM) biosensors that display up to a 20-fold fluorescence turn-on in response to binding hRAS (G12V variant), c-Abl Src homology 2 (SH2) domain, and WD repeat-containing protein 5 (WDR5) in cells, and >100-fold change in vitro. These sensors were created by inserting the previously developed hRAS^18,19^, SH2^20,21^, and WDR5^22^-binding MBs into mTurquoise and YFP from *Aequorea victoria* and mTag-RFP from *Entacmaea quadricolor*, at three different loops within the FPs. Unlike simple reporter molecules whose fluorescence is unchanged in bound and free states, ATOM sensors turn on because of binding. We demonstrated this capability in live and fixed human cells by detecting hRAS, SH2, and WDR5 proteins that were localized to the plasma membrane, cytoplasm, and nucleus (respectively) in the same cells, using multiplexed tri-color imaging, and sensing ER and mitochondria-resident SH2 protein using biosensor and SH2 that were targeted to those organelles. The ATOM turn-on mechanism is distinct from that of GECIs and involves ligand-assisted chromophore maturation. This work defines a method for the high-probability discovery of additional ATOM sensors by fusing new MBs, circularly permuted at one of two locations, into one of three surface loops in the FP. This six-member library can be rapidly screened in cells for the biosensor that yields the brightest turn-on in response to the co-expressed ligand.

## Results

### Biosensor design and optimization

The sensor input domains consisted of MBs (∼100 AA) that had been previously engineered to recognize SH2^20^, WDR5^22^, and hRAS^19^. To make the distance between their amino and carboxy termini close enough to be compatible for insertion into surface turns of the FPs, we circularly permuted the MBs at one of two loops that connect its seven β-strands (strands A – G; Figure 1A). The two MB variants are denoted MB1 (permuted in the CD loop) and MB2 (permuted in the EF loop), with superscripted SH2, WDR5, or RAS to indicate the binding specificities of the parental MBs. The CD and EF loops were chosen because they lie on the side of the molecule opposite to that of the ligand binding interface (generally the BC and FG loops). Some ‘side binding’ ligands such as hRAS interact with residues in strand D, which prompted us to make MB2 in which the permutation site is farther removed from the ligand binding site. The domain arrangements of ATOM sensors used in this study are shown in Table 1.

**Figure 1.**
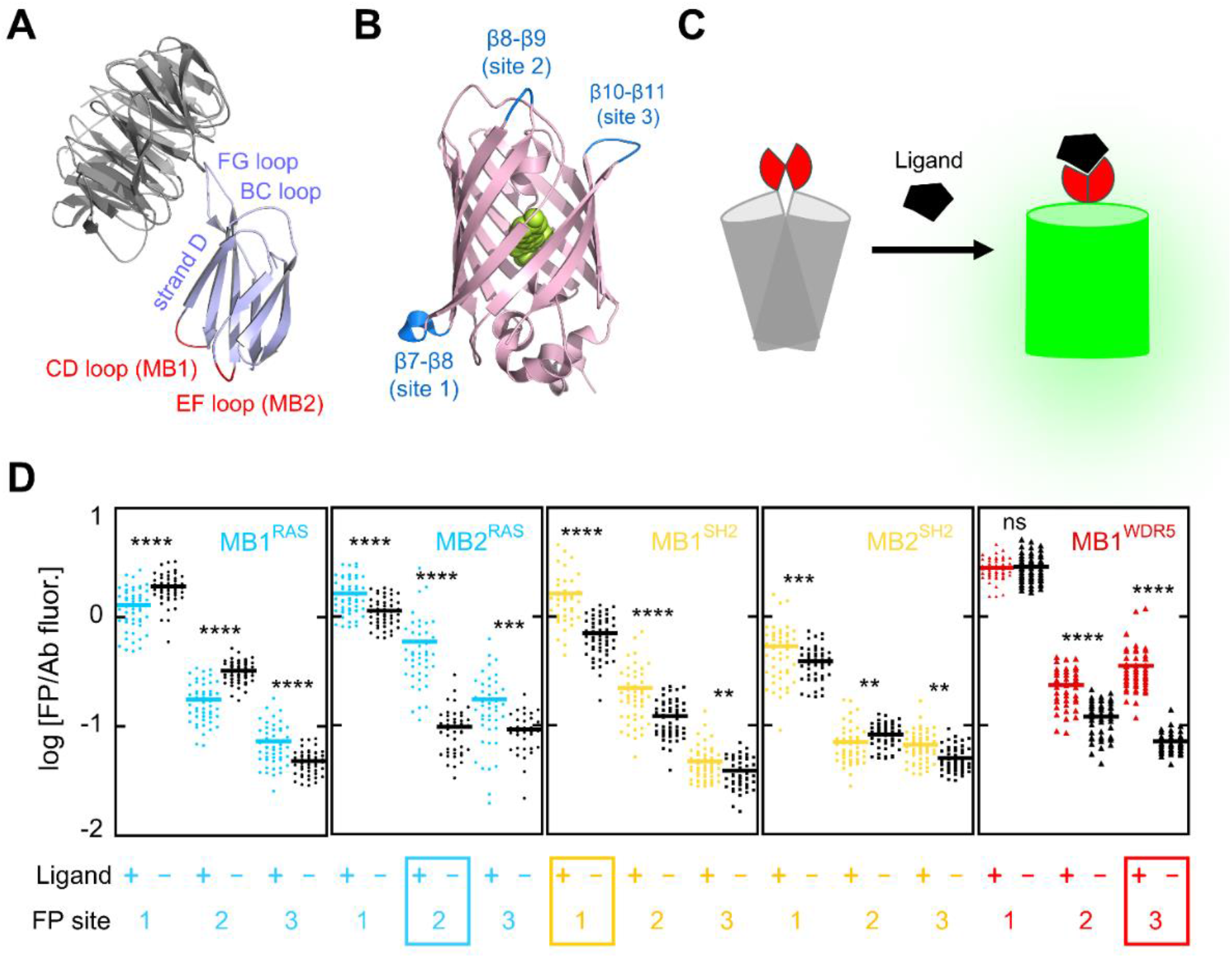
Biosensor design and screening y-ATOM sensors in HEK 293T cells. **(A)** The X-ray structure of MB^WDR5^ (purple) bound to WDR5 (grey) is shown, with loops in which MB1 (CD loop) and MB2 (EF loop) were permuted colored red. PDB 6BYN. **(B)** MB1 or MB2 were inserted into YFP (pink) at one of the three sites corresponding to the loops depicted in blue. The YFP chromophore is shown in green spheres. PDB 5WJ2. **(C)** In the proposed turn-on mechanism, the MB1/MB2 domains (red) are partially unfolded in the absence of ligand (black), which disrupts the conformation of the FP domain (grey), leaving the chromophore in an unmatured state. Ligand binding induces MB1/MB2 to fold, restoring the native conformation of the FP and allowing the chromophore to mature. **(D)** Fluorescent turn-on of y-ATOM sensors was determined in HEK 293T cells, with the best-performing y-ATOM sensor for each ligand indicated by the colored boxes. Cognate and noncognate ligands for each sensor are designated by plus and minus symbols, respectively. Each data point represents a cell. Significance was determined by a *t*-test with unequal variance: ****, p<10^−4^; ***, p<10^−3^; **, p<10^−2^; ns, not statistical. The results are representative of at least four biological repeats. Sample sizes and raw data are in Supporting Table S1.

**Table 1.**
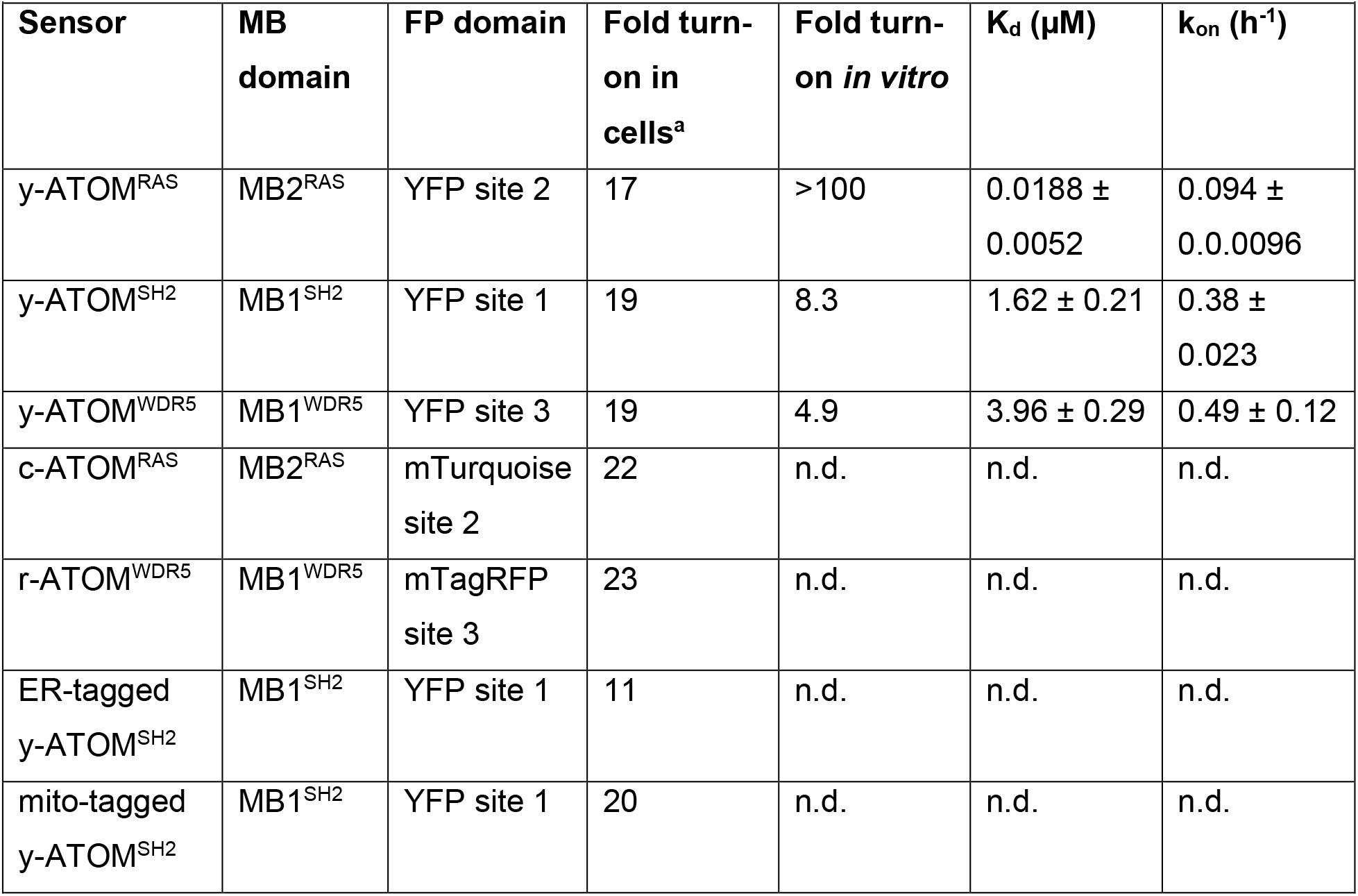
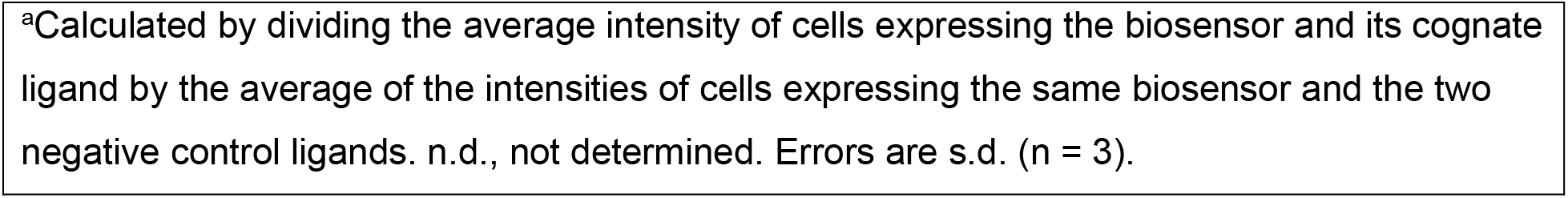
Properties of ATOM sensors.

| **Table 1. Properties of ATOM sensors.** |  |  |  |  |  |  |
| --- | --- | --- | --- | --- | --- | --- |
| **Sensor** | **MB domain** | **FP domain** | **Fold turn-on in cells<sup>a</sup>** | **Fold turn-on *in vitro*** | **K<sub>d</sub> ($\mu\text{M}$)** | **k<sub>on</sub> (h<sup>-1</sup>)** |
| y-ATOM<sup>RAS</sup> | MB2<sup>RAS</sup> | YFP site 2 | 17 | >100 | 0.0188 ± 0.0052 | 0.094 ± 0.0096 |
| y-ATOM<sup>SH2</sup> | MB1<sup>SH2</sup> | YFP site 1 | 19 | 8.3 | 1.62 ± 0.21 | 0.38 ± 0.023 |
| y-ATOM<sup>WDR5</sup> | MB1<sup>WDR5</sup> | YFP site 3 | 19 | 4.9 | 3.96 ± 0.29 | 0.49 ± 0.12 |
| c-ATOM<sup>RAS</sup> | MB2<sup>RAS</sup> | mTurquoise site 2 | 22 | n.d. | n.d. | n.d. |
| r-ATOM<sup>WDR5</sup> | MB1<sup>WDR5</sup> | mTagRFP site 3 | 23 | n.d. | n.d. | n.d. |
| ER-tagged y-ATOM<sup>SH2</sup> | MB1<sup>SH2</sup> | YFP site 1 | 11 | n.d. | n.d. | n.d. |
| mito-tagged y-ATOM<sup>SH2</sup> | MB1<sup>SH2</sup> | YFP site 1 | 20 | n.d. | n.d. | n.d. |

<sup>a</sup>Calculated by dividing the average intensity of cells expressing the biosensor and its cognate ligand by the average of the intensities of cells expressing the same biosensor and the two negative control ligands. n.d., not determined. Errors are s.d. (n = 3).

For initial biosensor development, we chose for the output domain a yellow variant of Clover FP (YFP) that had been circularly permuted between strands β9 and β10 of the 11-stranded β-barrel (Figure 1B)^17,23^. We previously found that inserting a circularly permuted FK506 binding protein in the β10-β11 loop (site 3 in Figure 1B) of the above YFP construct resulted in ligand-dependent turn on, with the proposed mechanism shown schematically in Figure 1C^17^. For the present study we inserted MB1 or MB2 into site 3 and explored two other FP loops as well: β7-β8 (site 1) and β8-β9 (site 2).

We screened the library of y-ATOM biosensors in human cells to determine which combinations of FP fusion site and MB permutation site displayed the greatest fluorescent turn-on in the presence of ligand. For each ligand there were six possible y-ATOM biosensors; we tested 15 of the 18 possible combinations for all three ligands (Figure 1D and Supporting Figures S1 – S3). HEK 293T cells were co-transfected with one plasmid encoding the sensor and a second plasmid expressing the target ligand or one of the two non-cognate ligands. The cells were then stained with an anti-GFP antibody to verify biosensor expression, and the signal from each sensor was quantified by dividing the FP intensity by the antibody staining intensity. Most MB/FP combinations exhibited significant turn-on (Figure 1D). The best performing yellow sensor for each ligand (y-ATOM^RAS^, y-ATOM^SH2^, and y-ATOM^WDR5^) are shown in cyan, yellow, and red boxes in Figure 1D. Curiously, no single combination of YFP fusion location and MB permutation site proved to be the most successful. The YFP insertion sites in y-ATOM^SH2^, y-ATOM^RAS^, and y-ATOM^WDR5^ were site 1, site 2, and site 3, respectively. MB1 gave the best results for y-ATOM^WDR5^ and y-ATOM^SH2^, while MB2 worked optimally for y-ATOM^RAS^.

### Biosensor specificity in cells

Having determined the optimal combination of MB permutation and FP insertion for each y-ATOM sensor, we next evaluated their ability to discriminate between cognate and decoy ligands in cells. HEK 293T cells were co-transfected with two plasmids: one harboring the y-ATOM^RAS^, y-ATOM^SH2^, or y-ATOM^WDR5^ gene, and the other expressing hRAS, SH2, or WDR5. All three y-ATOM sensors demonstrated high turn-on for their specific ligands (17 – 20-fold) compared to the two negative control ligands (Figure 2 and Table 1).

**Figure 2.**
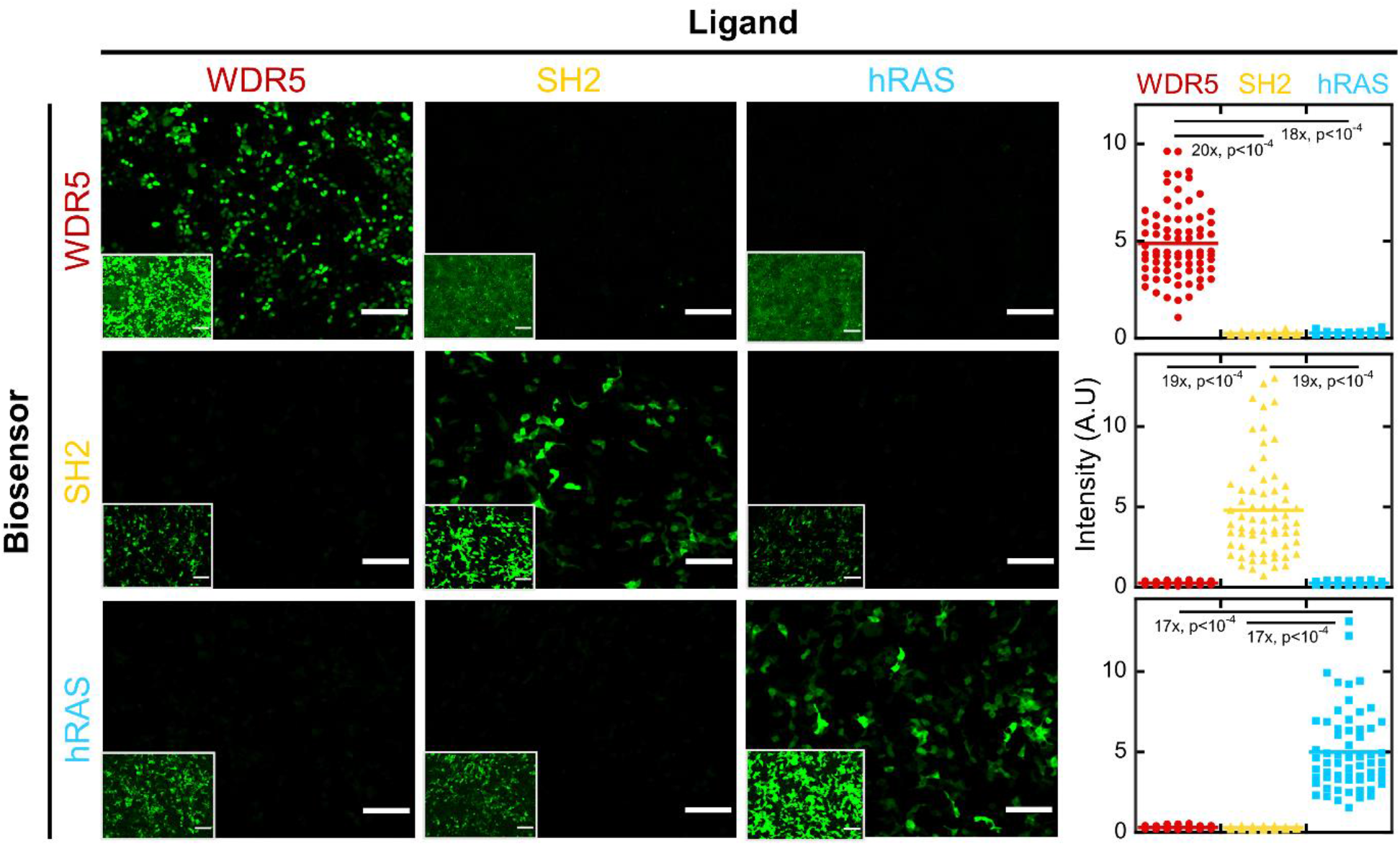
y-ATOM sensors show high selectivity and fluorescence turn-on in human cells. Images on the diagonal represent cells transfected with plasmids expressing biosensor and cognate ligand; off-diagonal images are cells transfected with plasmids expressing biosensor and decoy ligands. Higher contrast settings are shown in the insets to demonstrate that biosensors were expressed (but dim) in the off-diagonal cells. Fluorescence turn-on is quantified at right. Scale bars are 100 µm. Significance was determined by a *t*-test with unequal variance. The results are representative of four biological repeats. Sample sizes and raw data are in Supporting Table S1.

To establish whether the observed response resulted from biosensor turn-on or from an increase in biosensor concentration (e.g., due to decreased turnover and/or increased synthesis), we transfected cells with y-ATOM sensors and ligands as above, fixed them after 48 h, and then stained the cells with an anti-GFP antibody conjugated with the red fluorescent dye Alexa594. Red fluorescence remained constant in cells expressing hRAS, SH2, and WDR5 (Supporting Figure S4), indicating that biosensor levels were similar in the presence of target ligand and decoy ligands. Turn-on was therefore likely due to increased brightness of the sensor proteins. Supporting Figure S4 also demonstrates that the biosensors functioned in fixed cells as well as live cells.

### Biophysical characterization of sensors

To gain further insight into mechanism and to determine binding affinities and kinetics, we expressed y-ATOM^RAS^, y-ATOM^SH2^, and y-ATOM^WDR5^ in *E. coli* at 37 °C and purified the proteins using nickel-NTA chromatography. As anticipated, the sensors were mostly non-fluorescent (Figure 3A). This was due to the absence of the YFP chromophore, which could be identified by an absorbance peak at 514 nm. There was no evidence of mature (but protonated) chromophore as this species absorbs near 400 nm^2,24^. After the ligands were added, the 514 nm absorbance peaks appeared as did the expected fluorescence peak at 525 nm (Figure 3A).

**Figure 3.**
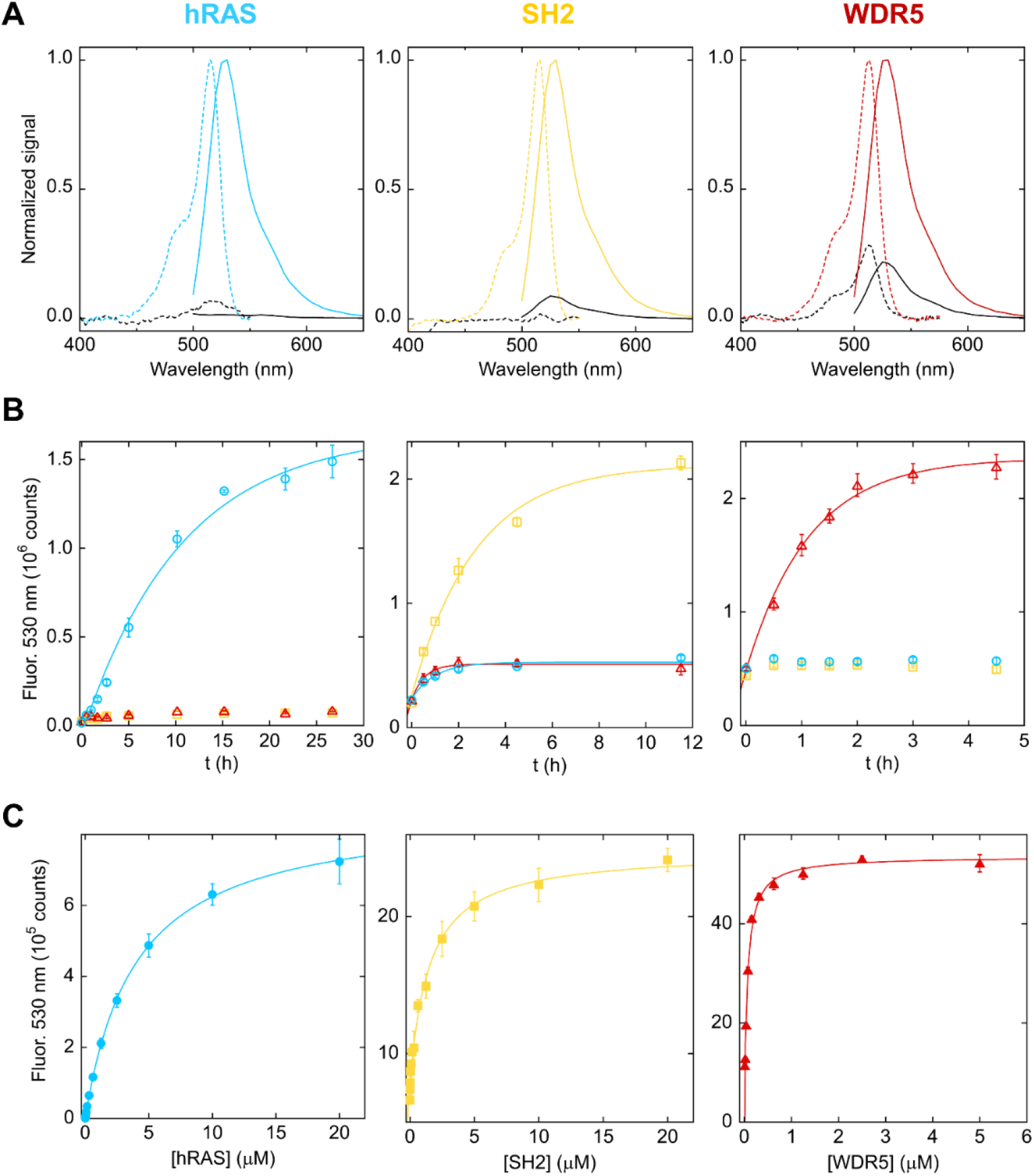
In vitro characterization of y-ATOM sensors at 37 °C. Sensors and ligands are colored cyan (hRAS), yellow (SH2), and red (WDR5). **(A)** Fluorescence turn-on resulted from ligand-induced chromophore maturation, as evidenced by the increase in absorbance at 514 nm (dashed lines) and fluorescence at 530 nm (solid lines). Samples contained 20 µM ligand. Data were recorded at time zero (dashed lines) and 24 h (solid lines). **(B)** Fluorescence was activated in a ligand-specific manner with half-times of 1.7 – 7.4 h. Lines are best fits of the data to a single-exponential function. **(C)** y-ATOM sensors bound their cognate targets with low nM to low µM K_d_. Lines are best fits of the data to the one-site binding equation. Error bars are s.d. (n = 3 technical repeats; 2 – 3 biological repeats were performed with similar results).

Fitting the time-dependent increases in fluorescence to single exponential functions revealed similar turn-on rates for y-ATOM^WDR5^ (k_on_ = 0.49 ± 0.12 h^-1^) and y-ATOM^SH2^ (k_on_ = 0.38 ± 0.023 h^-1^) at 37 °C (Figure 3B and Table 1). These values are somewhat slower than the maturation rate of Clover FP (1.9 h^-1^)^25^, although the maturation rate of the circularly permuted, yellow variant (Tyr203) of Clover that we used here has not been determined. Nevertheless, the data suggest that k_on_ may be partially limited by a binding-induced conformational change step that precedes the chromophore maturation reaction. This scenario is almost certainly present for y-ATOM^RAS^, which turned on ∼4-fold more slowly (k_on_ = 0.094 ± 0.0096 h^-1^) than y-ATOM^WDR5^ and y-ATOM^SH2^.

The fluorescent turn-on ratios of the y-ATOM sensors were calculated by dividing the fluorescence at the longest time point by the fluorescence at time zero, from the data in Figure 3B. Turn-on ratios were >100-fold (y-ATOM^RAS^), 4.9-fold (y-ATOM^WDR5^), and 8.3-fold (y-ATOM^SH2^) (Table 1). It is likely that the lower values observed for y-ATOM^WDR5^ and y-ATOM^SH2^ in vitro compared to in cells (17 – 20-fold; Figure 2) were due to ligand-independent turn-on that occurred during protein purification (∼3 h at room temperature). At this lower temperature we observed slow chromophore maturation in the absence of ligands, presumably because the MB and/or FP domains were more stable than at 37 °C. y-ATOM^RAS^ did not mature appreciably during purification, most likely because its intrinsic turn-on rate was ∼4-fold slower than those of y-ATOM^WDR5^ and y-ATOM^SH2^.

Turn-on of y-ATOM^WDR5^ and y-ATOM^RAS^ were almost completely dependent on their respective ligands, but y-ATOM^SH2^ showed a minor increase in fluorescence in the presence of the negative control ligands, indicating a nonspecific activation process (Figure 3B). The nonspecific activation rates (1.0 – 1.8 h^-1^) were slightly faster than the specific activation rate for reasons that are not clear. To test for reversibility, we added excess competitor (MB1^WDR5^) to the complex of y-ATOM^WDR5^ and WDR5. No decrease in fluorescence was observed, indicating that activation is irreversible (Supplemental Figure S5). This result is consistent with a chromophore maturation mechanism.

We next determined binding affinities by monitoring the increase in YFP fluorescence as a function of ligand concentration after 24 h of incubation at 37 °C. Fitted dissociation constants (K_d_) were 18.8 ± 5.2 nM (y-ATOM^WDR5^), 1.62 ± 0.21 µM (y-ATOM^SH2^), and 3.96 ± 0.29 µM (y-ATOM^RAS^) (Figure 3C and Table 1). These K_d_ values are higher than those of the parental (non-circularly permuted) MBs by factors of 3.8 (WDR5)^22^, 74 (SH2)^20^, and 240 (hRAS)^19^. It is likely the loss of affinity was due to a combination of circular permutation, coupling of ligand binding energy to conformational changes in the MB and FP domains, and possible steric effects of fusion to the FP protein.

### Multiplexed cellular imaging using three-color ATOM sensors

We next designed ATOM sensors of two additional colors, cyan and red. mTurquoise differs from Clover by only seven residues, so we created the cyan sensor c-ATOM^RAS^ by circularly permuting mTurquoise at the same position we did for Clover and then inserting MB2^RAS^ into the same loop (site 2) that gave the best results for y-ATOM^RAS^. We turned to the red FP scaffold mTagRFP to generate r-ATOM^WDR5^. We additionally used the unmodified mTagRFP sequence to determine whether ATOM sensors could be created without first circularly permuting the FP. Because of this difference, and because mTagRFP and Clover have very low sequence identity and possess different biophysical and photophysical properties, we repeated the optimization process by fusing MB1^WDR5^ into the three surface loops of mTagRFP that corresponded to those shown in Figure 1B. Each construct was then transiently co-transfected with WDR5 or SH2 expressing plasmids to quantify signal and background, respectively. The insertion point that yielded the highest turn-on normalized by total sensor expression was site 3, the same location that produced the greatest turn-on for y-ATOM^WDR5^ (Supporting Figure S6).

Specificity of the three-colored sensors was evaluated by co-transfecting HEK 293T cells with an equimolar mixture of r-ATOM^WDR5^, c-ATOM^RAS^, and y-ATOM^SH2^ plasmids and a fourth plasmid expressing either WDR5, hRAS, or SH2. WDR5, hRAS, and SH2 localize primarily to the nucleus^26^, cytosolic face of plasma membrane^19^, and cytoplasm^27^, respectively. We therefore fused a nuclear localization signal (NLS) to r-ATOM^WDR5^ and a nuclear export signal (NES) to c-ATOM^RAS^. The purpose of the NES was to reduce background signal coming from the small amount of ligand and sensor that entered the nucleus due to inefficient ribosome skipping of the polycistronic expression vector that encoded for the three ligands (*vide infra)*. This occurred with all sensors, but even weak nuclear fluorescence can obscure visualization of plasma membrane proteins^28^. y-ATOM^SH2^ was not tagged with an NES or NLS. As expected, we observed significant levels of only red fluorescence with WDR5 ligand (23-fold turn-on), only yellow fluorescence with SH2 ligand (19-fold turn-on), and only cyan fluorescence with hRAS ligand (22-fold turn-on) (Supporting Figure S7 and Table 1).

We next asked whether the red, cyan, and yellow ATOM sensors could detect all three targets in the same cell, at their proper locations, in a multiplexed experiment. The three-color biosensor plasmid mix was co-transfected with fourth plasmid harboring hRAS, SH2, and WDR5 genes separated by the P2A ribosome skipping sequence. All three biosensors showed robust activation when co-transfected with the plasmid expressing the ligands (Figure 4A, top and middle rows). Quantification revealed turn-on ratios of 19-fold (cyan channel), 13-fold (yellow channel), and 6-fold (red channel) (Figure 4B). Tracing a line through a single cell indicated sharp boundaries between nuclear, cytoplasmic and plasma membrane compartments and confirmed the expected localizations of hRAS, SH2, and WDR5, respectively (Figure 4C).

**Figure 4.**
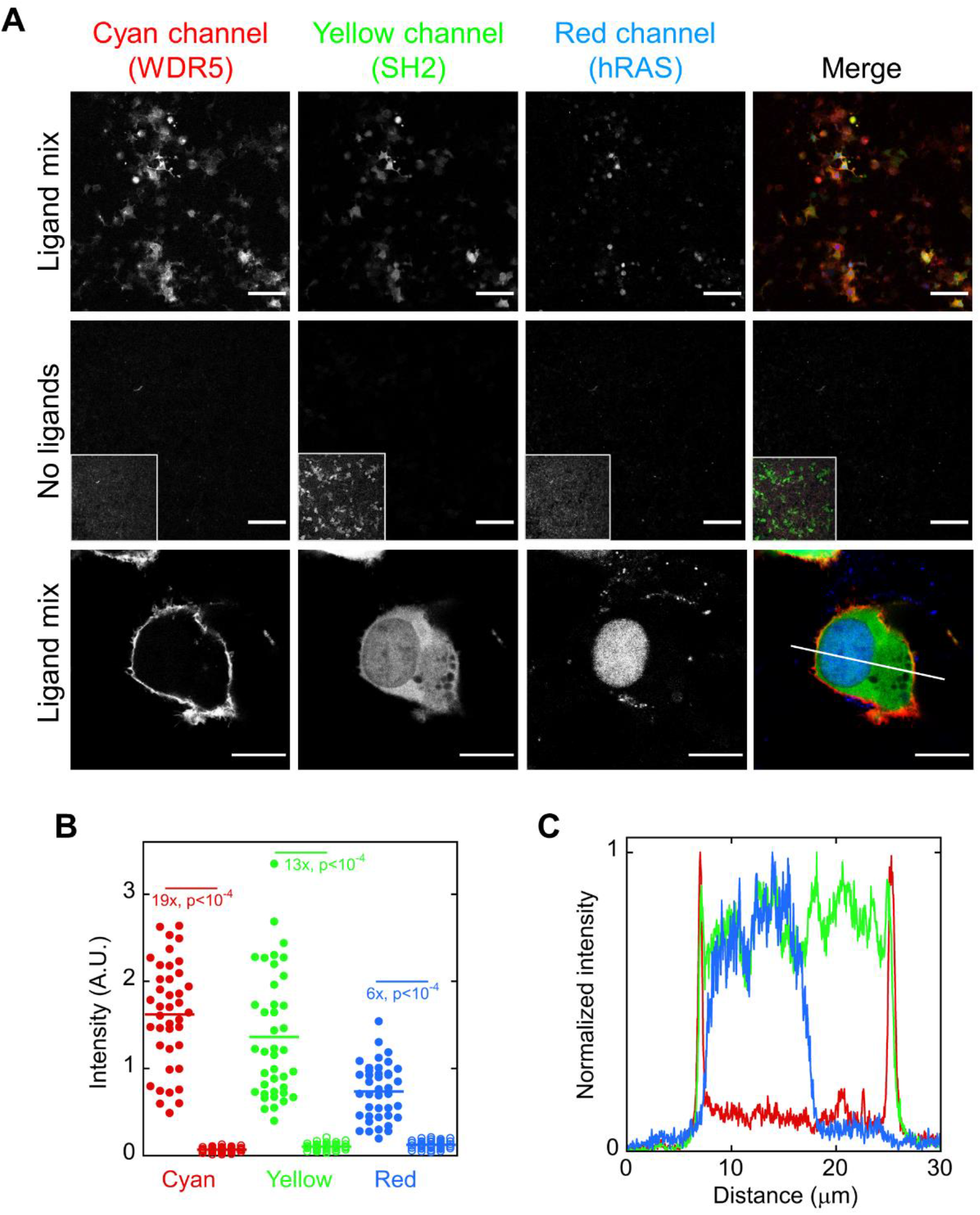
Multiplexed 3-color imaging shows ATOM sensors detect subcellular localization of protein targets. For visual clarity the cyan channel is represented in red, the yellow channel in green, and the red channel in blue. **(A)** Cells co-transfected with a mixture of c-ATOM^RAS^, y-ATOM^SH2^, r-ATOM^WDR5^ plasmids and a fourth plasmid encoding all three ligands showed fluorescent turn-on specific to the presence of ligands (top row and middle row). The higher contrast settings in the insets demonstrated that biosensors expressed but were dim. Focusing on a single cell revealed distinct biosensor fluorescence localized to the plasma membrane (c-ATOM^RAS^), cytoplasm (y-ATOM^SH2^), and nucleus (r-ATOM^WDR5^). Scale bars are 100 µm in top and middle rows, and 10 µm in the bottom row. **(B)** Sensor turn-on was 19-fold for hRAS (cyan channel), 13-fold for SH2 (yellow channel), and 6-fold for WDR5 (red channel). Filled and empty circles represent individual cells from the top and middle rows of panel A, respectively. Significance was determined by a *t*-test with unequal variance. The results are representative of three biological repeats. Sample sizes and raw data are in Supporting Table S1. **(C)** A line trace through the cell shown in the bottom row of panel A confirmed the expected locations of hRAS, SH2, and WDR5 in the plasma membrane, cytoplasm, and nucleus, respectively.

The above results demonstrate that both *Aequorea victoria* and *Entacmaea quadricolor* FPs can serve as output domains for the ATOM sensor mechanism. The *A. victoria* lineage covers blue to yellow FPs, and the *E. quadricolor* lineage extends emission from orange to far-red. The ATOM family of sensors therefore has the potential to span most of the visible spectrum, potentially allowing the detection of four or more colors in the same cell.

### Detection of proteins in mitochondria and endoplasmic reticulum

Mitochondria and ER can be challenging environments for protein-based biosensors, as the molecules must unfold and refold as they are translocated across the membranes, and the oxidative conditions in the ER can interfere with function of some FPs^29^. To establish whether ATOM sensors could be used for detecting targets in mitochondria and ER, we tagged y-ATOM^SH2^ and SH2 with peptides that directed them to those compartments. HeLa cells were co-transfected with a plasmid expressing ER-tagged biosensor and a second plasmid encoding ER-tagged SH2, mitochondrial (mito)-tagged SH2, or untagged SH2 (which expresses in the cytoplasm [Figure 4]). The ER-tagged biosensor exhibited very little fluorescence in cells expressing mito-tagged SH2 and untagged SH2 (Figure 5A), and strong fluorescence in cells expressing ER-tagged SH2. Similarly, cells transfected with mito-tagged biosensor were mostly dark in the cells expressing ER-tagged SH2 and untagged SH2 and were bright in cells expressing mito-tagged SH2 (Figure 5A). Quantification found turn-on ratios of 9 – 13-fold for ER-tagged biosensor/ligand and 19 – 21-fold for mito-tagged biosensor/ligand, relative to the negative controls (Figure 5C).

**Figure 5.**
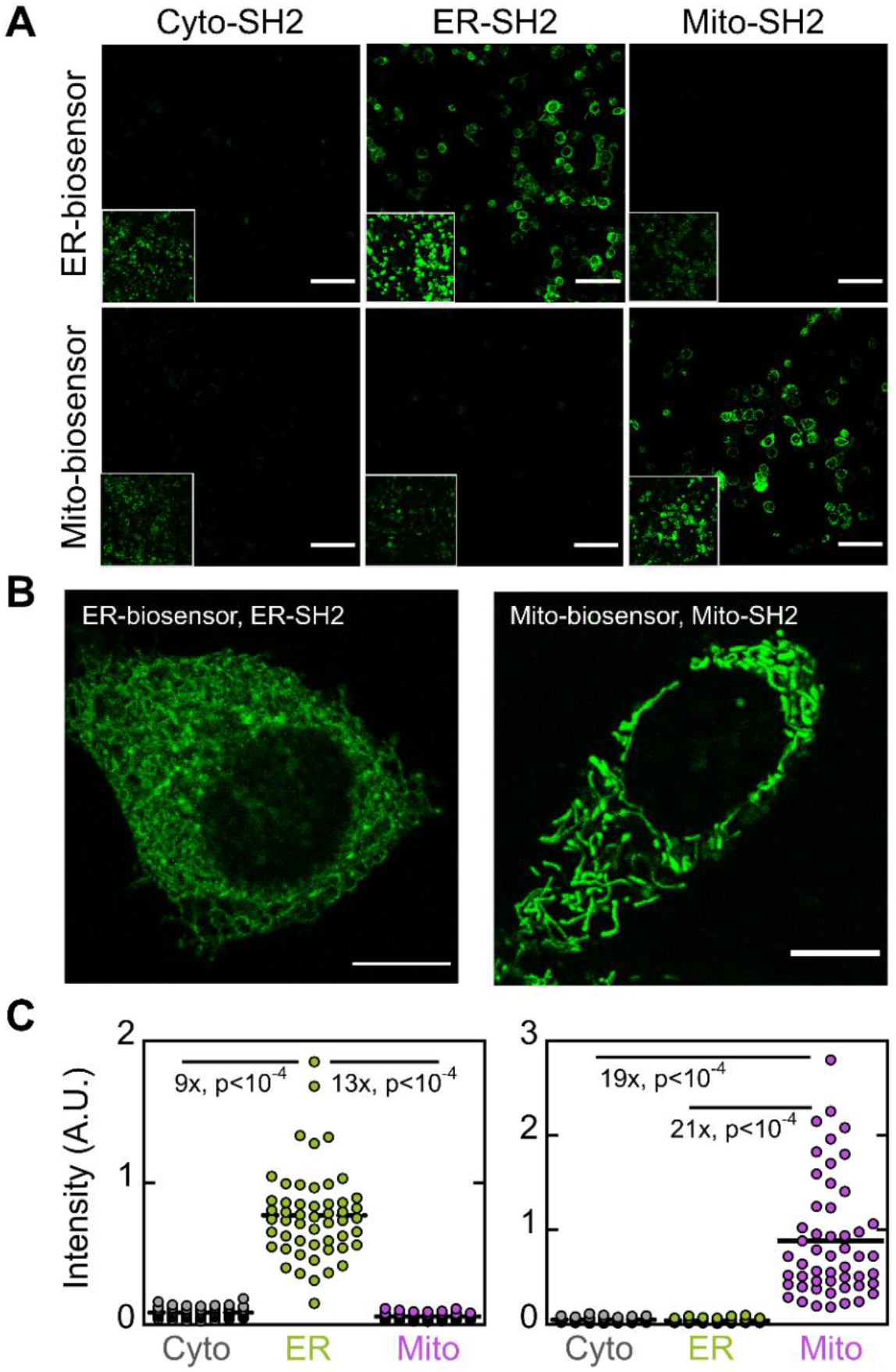
y-ATOM^SH2^ can function as an ER- or mitochondria-targeted biosensor. **(A)** HeLa cells transfected with ER-tagged biosensor (top row) or mito-tagged biosensor (bottom row) were activated specifically by ER-tagged SH2 and mito-tagged SH2, respectively, and not by SH2 targeted other compartments. Insets show higher contrast settings to demonstrate that biosensors were expressed in all transfection conditions. Scale bars are 100 µm. **(B)** Confocal images confirmed that the biosensors were detecting their targets in the expected organelles. Scale bars are 10 μm. **(C)** Quantification of the data in panel A found 9 – 13-fold selective turn-on for ER-targeted sensor and ligand, and 19 – 21-fold selective turn-on for mito-targeted sensor and ligand. Significance was determined by a *t*-test with unequal significance. The results are representative of three biological repeats. Sample sizes and raw data are in Supporting Table S1.

Confocal images confirmed that the biosensors were detecting their targets in the correct organelles, showing a netlike pattern of the ER and the tubular structure of mitochondria (Fig. 5B). These results demonstrate that ATOM biosensors remain functional after translocation into the ER and mitochondria. Of note, the relatively slow k_on_ rate is advantageous for this application. If that rate were fast, the sensors may become activated in the cytoplasm if their targets are cytosolic and remain fluorescent after transport into ER/mitochondria, causing nonspecific background in those organelles. Figure 5A demonstrates that this false positive mechanism does not occur appreciably with y-ATOM^SH2^.

## Discussion

The ATOM biosensors combine a fluorescent reporter protein with a MB input domain to detect binding of target ligands with high specificity. Because they are ON/OFF switches, multiple ATOM sensors of different colors can be simultaneously overexpressed in cells, whereupon they become fluorescent only when encountering their ligands in their specific cellular locations. This property was demonstrated by detecting three separate proteins in a single mammalian cell, in locales including plasma membrane, cytoplasm, nucleus, endoplasmic reticulum, and mitochondria. This result would not have been feasible using simple, overexpressed reporter proteins, e.g., MBs or other binding domains fused at one of their termini to GFP.

ATOM sensors differ from the GECIs in several notable respects, and each has its strengths and weaknesses. First, the ATOM sensing mechanism involves irreversible chromophore maturation, while that of GECIs entails reversible deprotonation of the chromophore. GECIs therefore excel at reporting on rapid (ms scale) fluctuations in analyte concentration, whereas ATOM sensors are best suited to detect intracellular targets that may be present at low abundance and when signal accumulation is desired or acceptable. The chromophore maturation mechanism lends itself to high sensitivity, as unmature chromophores are completely dark and thus produce large signal-to-noise ratios upon maturation. The second distinction is that ATOM sensors are designed from the ground up to be adaptable by virtue of the MB domain. MBs can be evolved *in vitro* to recognize new ligands, and there is no requirement for a binding-induced conformational change. By contrast, each GECI-type biosensor is constructed using a unique domain that typically dimerizes upon ligand binding (e.g., CaM/RS20 peptide), and this type of conformational change is not readily transferable to other targets.

There are at least two other sensing mechanisms that entail chromophore maturation. The first is bimolecular fluorescence complementation (BiFC), wherein a split FP is tagged to separate domains that dimerize upon ligand binding. BiFC is a robust mechanism, producing large turn-on ratios and typically irreversible activation, with its main limitation being the requirement that the recognition domain be split and able to reconstitute upon ligand binding^30^. The second mechanism, demonstrated by Hardy and colleagues^31,32^, involves inhibition of chromophore maturation when GFP is fused to a tetrameric peptide. They hypothesized that tetramerization caused GFP to adopt a strained but folded conformation in which the chromophore could not mature. Cleavage of the peptide relieved strain and allowed GFP to become fluorescent.

Because we observed ligand-assisted chromophore maturation with multiple combinations of MB permutation site and FP insertion loop, this phenomenon is not strictly dependent on a particular amino acid sequence or structure that comprises the interface between the two proteins. Moreover, since MBs recognize their targets via loops that are distant from the FP insertion site, and MBs do not undergo a conformational change upon binding, it seems logical that global stabilization or folding of the MB domain may be the event that triggers chromophore maturation. We previously showed that binding-induced folding of unstable, circularly permuted FK506 binding protein stabilized the FP into which it was inserted by a mechanism known as loop-closure entropy (LCE)^17^, and that LCE can be a particularly efficient means to effect allosteric communication between domains^33^. If the LCE switching mechanism is operational in the present case, it is reasonable to speculate that new families of ATOM sensors may be created using altogether different binding domains (such as nanobodies and affibodies) that are circularly permuted and made unstable prior to inserting into the FP.

## Conclusions

Though the ATOM sensing mechanism remains to be fully delineated, we showed that it worked with three out of three arbitrarily chosen MBs, as well as FPs from jellyfish and anemone lineages. This result suggests that ATOM sensors with different binding specificities and colors can be generated rapidly from new MBs, with a minimum of optimization. To do so, we recommend synthesizing the six-member gene library described in Figure 1 and screening by co-transfection as shown in Figure 2. If our results are representative, at least one of the proteins will likely be a functioning ATOM sensor. This process stands in contrast to the large libraries and multiple iterations of screening and/or selection that are employed when developing many other single-molecule fluorescent biosensors^34,34,35^.

## Methods

### Gene construction

For bacterial expression, C-terminal 8x-His tagged biosensor genes were cloned into pET41 expression vector downstream of the T7 promoter using NdeI/XhoI restriction sites. The y-ATOM^RAS^ sensor expressed poorly in *E. coli*, so ribose binding protein (RBP) was fused to its N-terminus to improve solubility and overall expression^36^. y-ATOM^WDR5^ and y-ATOM^SH2^ did not contain RBP fusions.

For mammalian cell expression, ATOM genes were cloned into the N1 vector (Clontech) under the CMV promoter using AgeI/NotI restriction sites. The genes were then fused to the 3’-end of the maltose binding protein (MBP) gene using the EcoRI/AgeI sites. WDR5, SH2, and hRAS (G12V variant) proteins were constructed without any exogenous localization tags. The C-terminal CAAX farnesylation sequence was deleted from hRAS in all experiments except for the multiplexed three-color detection study (Figure 4). For that latter experiment, the r-ATOM^WDR5^ and c-ATOM^Ras^ proteins were tagged with a nuclear localization signal (MPKKKRKV) and nuclear export signal (MNLVDLQKKLEELELDEQQG), respectively, at the N-terminus of MBP. The ER-localized y-ATOM^SH2^ sensor and ligand were tagged with the prolactin secretion signal (MNIKGSPWKGSLLLLLVSNLLLCQSV) at their N-termini and the ER retention signal (KDEL) at their C-termini. The mitochondria-localized y-ATOM^SH2^ sensor and ligands were tagged with the COX4 signal peptide (MLSLRQSIRFFKPATRTLCSSRYLL) at their N-termini. All genes were fully sequenced. Amino acid sequences of biosensors and ligands are provided in Supplementary Figure S8.

### *In vitro* equilibrium binding and kinetic experiments

ATOM sensors were expressed in *E. coli* BL21(DE3) cells and induced with 0.3 mM isopropyl β-D-thiogalactopyranoside for 3 – 4 h at 37 °C. WDR5, SH2, and hRAS ligand proteins were expressed as above except cells were induced at 18 °C for 12 – 16 h. All sensor and ligand proteins except for SH2 were purified by nickel-nitrilotriacetate chromatography according to the manufacturer’s instructions (Molecular Cloning Laboratories). SH2 did not have a His-Tag and was purified using an SP Sepharose column as described^37^. Sensors were purified at room temperature and ligands at 4 °C. Upon elution from the nickel nitrilotriacetate columns, ATOM sensors were desalted into experimental buffer (20 mM Tris, pH 8.0, 0.15 M NaCl, 0.005 % TWEEN-20) using DG10 columns (Bio-Rad) and experiments were begun immediately. Total purification time was ∼3 h.

Fluorescence kinetic experiments were performed by mixing 10 – 50 nM of biosensor with 20 µM of each ligand and transferring to a 37 °C bath at time zero. At the indicated times, 100 µl of samples were aliquoted onto a fully blackened 96-well plate (Corning CoStar) and fluorescence spectra were recorded using a Molecular Devices i3x plate reader (475 nm excitation, 500 – 650 nm emission). To measure binding affinities, a solution of biosensor (10 – 50 nM) plus ligand (20 µM SH2 or hRAS, or 5 µM WDR5) was serially diluted 2-fold into solutions of biosensor only to generate the indicated ligand concentrations. Samples were incubated for 24 h at 37 °C and fluorescence spectra were recorded as above.

### Cell culture and transfection

HEK293T and HeLa cells were cultured at 37 °C in DMEM containing 10 % FBS and 1x penicillin/streptomycin. Cells were split into a 12-well plate containing round 14 mm glass coverslips one day before transfection at a confluency of 60 – 80 %. HEK 293T cells were transfected using calcium phosphate method as previously described^17,33^. HeLa cells were transfected using the TransIT-2020 reagent (Mirus Bio) according to the manufacturer’s protocol. After 36 – 48 h of expression, cells were washed with and mounted on imaging media (DMEM, 25 mM HEPES, pH 7.0). All imaging was performed on live cells except the 3-color multiplexed imaging (Figure 3) and the antibody labeling experiments (Supporting Figures S1-S4 and S6), both of which were fixed with paraformaldehyde. Ligand:biosensor plasmid ratio was 1:5 for the screening experiments in Figure 1 in order to quantify weakly fluorescent biosensors; transfections in all other experiments contained equal ratios of ligand and biosensor plasmids.

### Antibody staining

After 36 – 48 h of expression, the cells were fixed with 2.5% paraformaldehyde in PBS for 20 min, followed by three 5 min washes in 20 mM Tris (pH 7.5), 150 mM NaCl. Permeabilization and blocking was then performed by incubating the cells in blocking buffer (5 % BSA, 0.25 % Triton X-100 in 20 mM Tris, pH 7.5, 150 mM NaCl) for 30 min. The rabbit anti-GFP-Alexa594 conjugate or rabbit-anti-RFP antibodies (ThermoFisher Scientific) were diluted 1:1000 in blocking buffer and the cells were labelled for 1 h at room temperature, followed by three 5 min washes with 20 mM Tris (pH 7.5), 150 mM NaCl. The anti-RFP antibodies were labelled with anti-rabbit-Alexa 488 conjugate at 1:1000 dilution in blocking buffer for 1 h at room temperature, followed by three 5 min washes with 20mM Tris (pH 7.5), 150 mM NaCl. The coverslips were then mounted in 50 % glycerol, 20 mM Tris (pH 8.5), 50 mM NaCl and sealed with nail polish.

### Imaging

Screening experiments (Figure 2, Supporting Figures S1-S4 and S6) employed a Zeiss Axioimager Z1 upright microscope with 10x/0.25 A-Plan objective. The high-resolution three color and ER/mitochondria localization experiments (Figures 4 and 5, respectively) were performed on a Leica SP8 inverted laser-scanning confocal microscope using 405 nm excitation for mTurqoise, 488 nm excitation for YFP and 552 nm excitation for mTagRFP. Low-magnification confocal images were obtained using a 20x dry objective with the pinhole opened to 5 Airy units, allowing imaging at a low laser power to avoid phototoxicity. The high-magnification images were taken using a 63x oil-immersion objective with pinhole closed to 1 Airy unit.

For each experiment, at least three fields containing at least 30 cells each were imaged, and all three fields were pooled for quantification of that sample. The images were quantified using Fiji^38^. Background was first subtracted using the built-in plugin using 50 pixels as the sliding ball radius and no further processing was applied. The intensity of each cell was measured by manually drawing a region of interest around each cell and determining the average intensity using the *Measure Particles* plugin.

### Reproducibility, statistics, and data availability

All experiments described in this study were performed a minimum of three times. The in-cell experiments consisted of >3 biological repeats with new cells freshly split and transfected on different days. For in vitro experiments, sensor proteins were purified at least 2 times for independent biological repeats, with 3 technical repeats of each preparation. The statistical significance, where stated, was determined by a t-test with unequal variance using Kaleidagraph 5.0 (Synergy Software). All raw and processed data and images generated in this paper will be posted on the Mendeley repository. Plasmids and other reagents will be made available upon request.

## Author contributions

Conceptualization, J.-H.H. and S.N.L.; methodology, H.S., J.-H.H., and S.N.L.; validation, formal analysis, and investigation, H.S., J.-H.H., M.F.P., S.B.P., P.O.M. and S.N.L.; writing – original draft, H.S. and S.N.L.; writing – review & editing, H.S., J.-H.H., and S.N.L..; visualization, H.S., J.-H.H., and S.N.L.; funding acquisition, H.S. and S.N.L.

## Acknowledgements

We thank Drs. Xiaowen Wang and Xin-Jie Chen for advice on mitochondrial imaging, Dr. Michael Cosgrove for the gift of the WDR5 expression plasmid, and Dr. Shohei Koide for the gift of the MB^SH2^ and SH2 genes. This work was supported by NIH grants F30 GM146428 to H.S. and R01 GM148448 to S.N.L.

## Literature cited

1. Nakai, J., Ohkura, M. & Imoto, K. A high signal-to-noise Ca2+ probe composed of a single green fluorescent protein. Nat. Biotechnol. 19, 137–141 2001.

2. Baird, G. S., Zacharias, D. A. & Tsien, R. Y. Circular permutation and receptor insertion within green fluorescent proteins. Proc. Natl. Acad. Sci. 96, 11241–11246 1999.

3. Nasu, Y., Shen, Y., Kramer, L. & Campbell, R. E. Structure- and mechanism-guided design of single fluorescent protein-based biosensors. Nat. Chem. Biol. 17, 509–518 2021.

4. van der Linden, F. H. et al.A turquoise fluorescence lifetime-based biosensor for quantitative imaging of intracellular calcium. Nat. Commun. 12, 7159 2021.

5. Inoue, M. et al.Rational design of a high-affinity, fast, red calcium indicator R-CaMP2. Nat. Methods 12, 64–70 2015.

6. Inoue, M. et al.Rational Engineering of XCaMPs, a Multicolor GECI Suite for In Vivo Imaging of Complex Brain Circuit Dynamics. Cell 177, 1346-1360.e24 2019.

7. Sun, X. R. et al.Fast GCaMPs for improved tracking of neuronal activity. Nat. Commun. 4, 2170 2013.

8. Chen, T.-W. et al.Ultrasensitive fluorescent proteins for imaging neuronal activity. Nature 499, 295–300 2013.

9. Binz, H. K., Amstutz, P. &Plückthun, A. Engineering novel binding proteins from nonimmunoglobulin domains. Nat. Biotechnol. 23, 1257–1268 2005.

10. Hamers-Casterman, C. et al.Naturally occurring antibodies devoid of light chains. Nature 363, 446–448 1993.

11. Stumpp, M. T., Binz, H. K. & Amstutz, P. DARPins: A new generation of protein therapeutics. Drug Discov. Today 13, 695–701 2008.

12. Amstutz, P. et al.Intracellular Kinase Inhibitors Selected from Combinatorial Libraries of Designed Ankyrin Repeat Proteins *. J. Biol. Chem. 280, 24715–24722 2005.

13. Nord, K. et al.Binding proteins selected from combinatorial libraries of an α-helical bacterial receptor domain. Nat. Biotechnol. 15, 772–777 1997.

14. Lajoie, M. J. et al.Designed protein logic to target cells with precise combinations of surface antigens. Science 369, 1637–1643 2020.

15. Langan, R. A. et al.De novo design of bioactive protein switches. Nature 572, 205–210 2019.

16. Koide, A., Bailey, C. W., Huang, X. & Koide, S. The fibronectin type III domain as a scaffold for novel binding proteins11Edited by J. Wells. J. Mol. Biol. 284, 1141–1151 1998.

17. John, A. M., Sekhon, H., Ha, J.-H. & Loh, S. N. Engineering a Fluorescent Protein Color Switch Using Entropy-Driven β-Strand Exchange. ACS Sens. 7, 263–271 2022.

18. Teng, K. W. et al.Selective and noncovalent targeting of RAS mutants for inhibition and degradation. Nat. Commun. 12, 2656 2021.

19. Spencer-Smith, R. et al.Inhibition of RAS function through targeting an allosteric regulatory site. Nat. Chem. Biol. 13, 62–68 2017.

20. Wojcik, J. et al.A potent and highly specific FN3 monobody inhibitor of the Abl SH2 domain. Nat. Struct. Mol. Biol. 17, 519–527 2010.

21. Koide, A., Wojcik, J., Gilbreth, R. N., Hoey, R. J. & Koide, S. Teaching an old scaffold new tricks: monobodies constructed using alternative surfaces of the FN3 scaffold. J. Mol. Biol. 415, 393–405 2012.

22. Gupta, A. et al.Facile target validation in an animal model with intracellularly expressed monobodies. Nat. Chem. Biol. 14, 895–900 2018.

23. Do, K. & Boxer, S. G. GFP Variants with Alternative β-Strands and Their Application as Light-driven Protease Sensors: A Tale of Two Tails. J. Am. Chem. Soc. 135, 10226–10229 2013.

24. Tsien, R. Y. The Green Fluorescent Protein. Annu. Rev. Biochem. 67, 509–544 1998.

25. Balleza, E., Kim, J. M. & Cluzel, P. Systematic characterization of maturation time of fluorescent proteins in living cells. Nat. Methods 15, 47–51 2018.

26. Wang, P. et al.WDR5 modulates cell motility and morphology and controls nuclear changes induced by a 3D environment. Proc. Natl. Acad. Sci. 115, 8581–8586 2018.

27. Sirvent, A., Benistant, C. & Roche, S. Cytoplasmic signalling by the c-Abl tyrosine kinase in normal and cancer cells. Biol. Cell 100, 617–631 2008.

28. High Cleavage Efficiency of a 2A Peptide Derived from Porcine Teschovirus-1 in Human Cell Lines, Zebrafish and Mice | PLOS ONE. https://journals.plos.org/plosone/article?id=10.1371/journal.pone.0018556.

29. Shariati, K. et al.A Superfolder Green Fluorescent Protein-Based Biosensor Allows Monitoring of Chloride in the Endoplasmic Reticulum. ACS Sens. 7, 2218–2224 2022.

30. Hu, C.-D., Chinenov, Y. & Kerppola, T. K. Visualization of Interactions among bZIP and Rel Family Proteins in Living Cells Using Bimolecular Fluorescence Complementation. Mol. Cell 9, 789–798 2002.

31. Nicholls, S. B. & Hardy, J. A. Structural basis of fluorescence quenching in caspase activatable-GFP. Protein Sci. Publ. Protein Soc. 22, 247–257 2013.

32. Nicholls, S. B., Chu, J., Abbruzzese, G., Tremblay, K. D. & Hardy, J. A. Mechanism of a genetically encoded dark-to-bright reporter for caspase activity. J. Biol. Chem. 286, 24977–24986 2011.

33. Sekhon, H., Ha, J.-H. & Loh, S. N. Enhancing response of a protein conformational switch by using two disordered ligand binding domains. Front. Mol. Biosci. 10, 2023.

34. Nasu, Y. et al.A genetically encoded fluorescent biosensor for extracellular L-lactate. Nat. Commun. 12, 7058 2021.

35. Wu, S.-Y. et al.A sensitive and specific genetically-encoded potassium ion biosensor for in vivo applications across the tree of life. PLOS Biol. 20, e3001772 2022.

36. Blanden, A. R. et al.Zinc shapes the folding landscape of p53 and establishes a pathway for reactivating structurally diverse cancer mutants. eLife 9, e61487 2020.

37. Zheng, H., Bi, J., Krendel, M. & Loh, S. N. Converting a Binding Protein into a Biosensing Conformational Switch Using Protein Fragment Exchange. Biochemistry 53, 5505–5514 2014.

38. Schindelin, J. et al.Fiji: an open-source platform for biological-image analysis. Nat. Methods 9, 676–682 2012.

