## Supplementary Information for "Adaptable, Turn-On Monobody (ATOM) Fluorescent Biosensors for Multiplexed Detection in Cells"

Harsimranjit Sekhon*, Jeung-Hoi Ha*, Maria F. Presti, Spencer B. Procopio, Paige O. Mirsky, Anna M. John, Stewart N. Loh

Department of Biochemistry and Molecular Biology, SUNY Upstate Medical University, Syracuse, NY, USA

**Contents**

Figures S1 – S7, related to Figures 1 – 4

Figure S8, containing amino acid sequences of sensors and ligands used in the study

Table S1, related to Figures 1, 2, 4 and 5

Table S2, related to Figures S4, S6, and S7

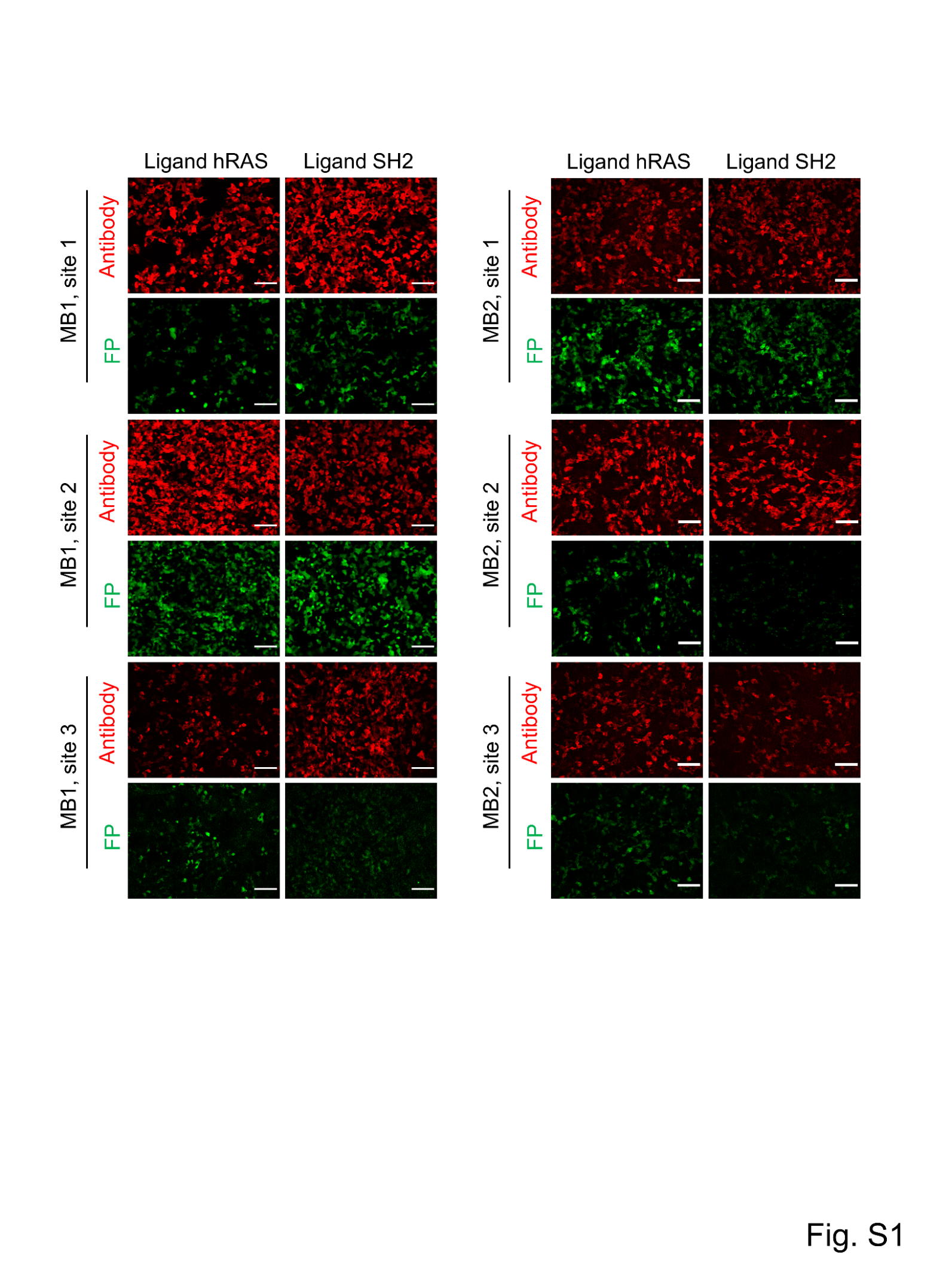

**Figure S1, related to Figure 1. Cell images showing screening of y-ATOM^RAS^ sensors.** y-ATOM^RAS^ sensors were composed of MB1^RAS^ or MB2^RAS^ fused to circularly permuted YFP at site 1, site 2, or site 3. HEK 293T cells were co-transfected with one plasmid encoding y-ATOM^RAS^ and a second plasmid expressing hRAS or SH2. Cells were fixed, stained with anti-GFP antibody and imaged as described in Methods. Scale bars are 100 µm. The results are representative of at least four biological repeats.

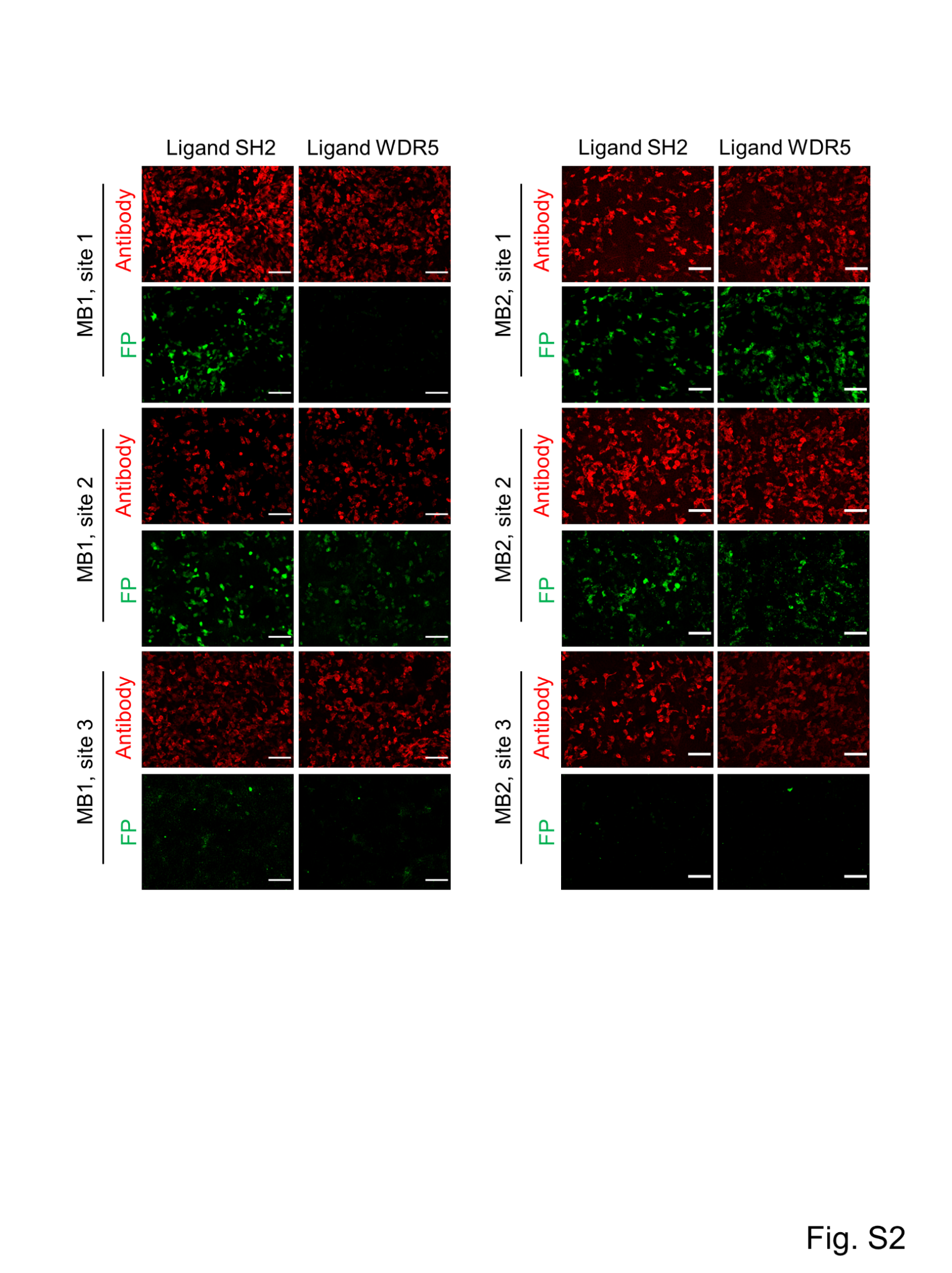

**Figure S2, related to Figure 1. Cell images showing screening of y-ATOM^SH2^ sensors.** y-ATOM^SH2^ sensors were composed of MB1^SH2^ or MB2^SH2^ fused to circularly permuted YFP at site 1, site 2, or site 3. HEK 293T cells were co-transfected with one plasmid encoding y-ATOM^SH2^ and a second plasmid expressing SH2 or WDR5. Cells were fixed, stained with anti-GFP antibody and imaged as described in Methods. Scale bars are 100 µm. The results are representative of at least four biological repeats.

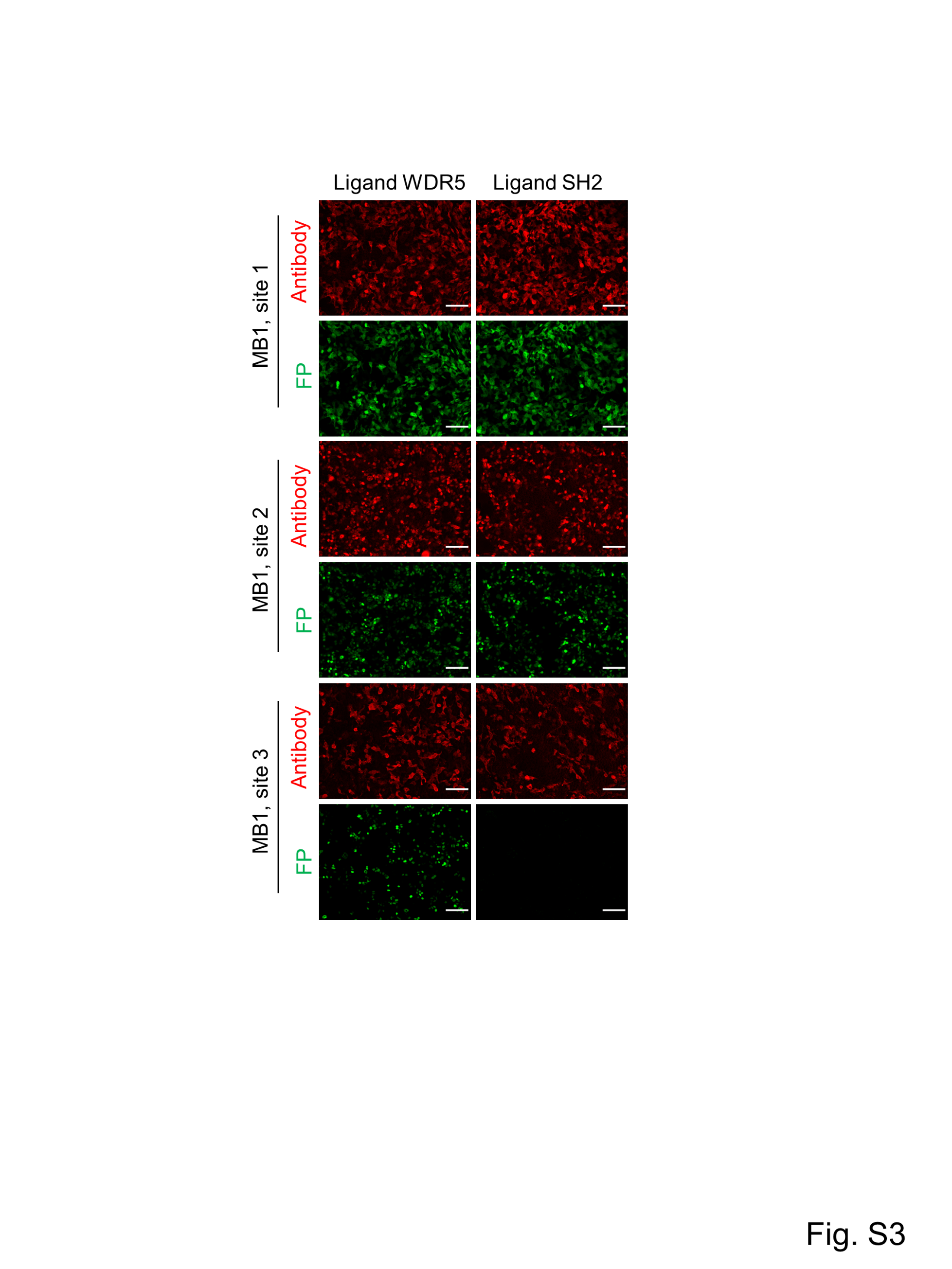

**Figure S3, related to Figure 1. Cell images showing screening of y-ATOM^WDR5^ sensors.** y-ATOM^WDR5^ sensors were composed of MB1^WDR5^ fused to circularly permuted YFP at site 1, site 2, or site 3. HEK 293T cells were co-transfected with one plasmid encoding y-ATOM^WDR5^ and a second plasmid expressing WDR5 or SH2. Cells were fixed, stained with anti-GFP antibody and imaged as described in Methods. Scale bars are 100 µm. The results are representative of at least four biological repeats.

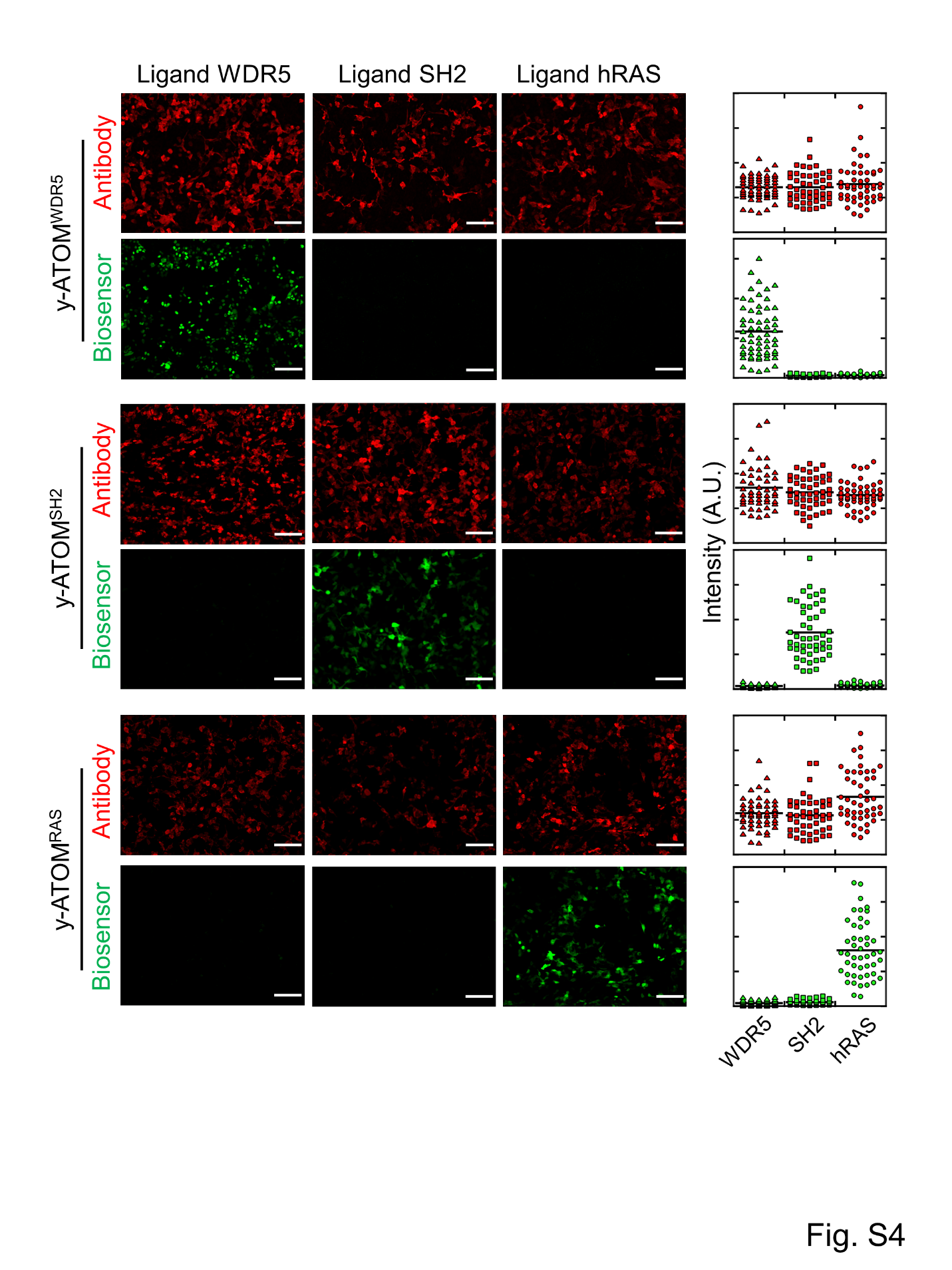

**Figure S4, related to Figure 2. Ligand-dependent ATOM turn-on in cells is not due to increased biosensor levels.** HEK 293T cells were co-transfected with one plasmid encoding y-ATOM^WDR5^, y-ATOM^SH2^, or y-ATOM^RAS^ and a second plasmid expressing WDR5, SH2, or hRAS. After 48 h, cells were fixed, stained with an anti-GFP antibody conjugated with Alexa594, and imaged in red (antibody) and yellow (biosensor; depicted as green) channels. Scale bars are 100 µm. Quantification is shown at right. Each data point represents one cell. The results are representative of three biological repeats.

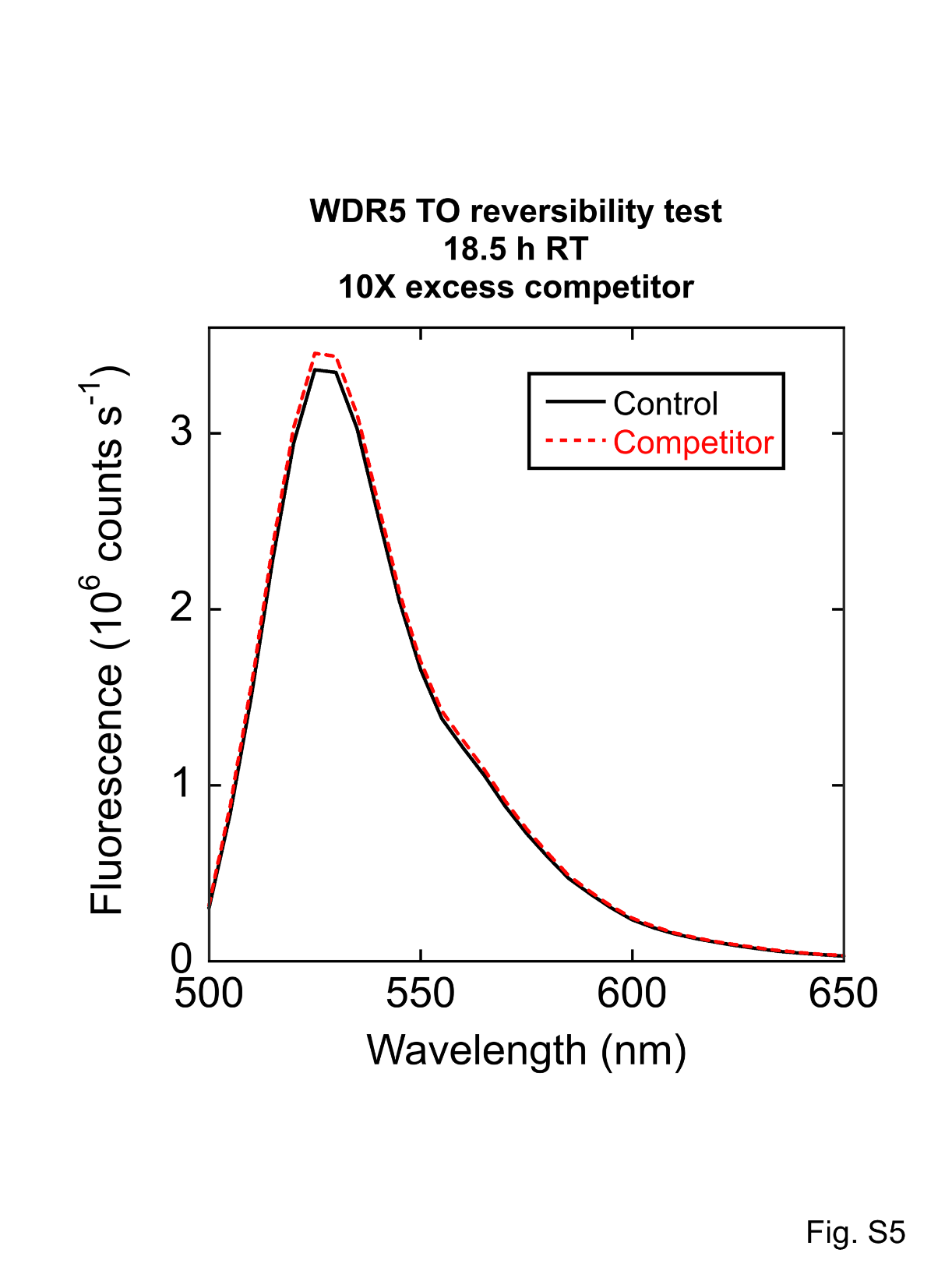

**Figure S5, related to Figure 3. Turn-on of ATOM biosensors is irreversible.** y-ATOM^WDR5^ was incubated with 2 µM WDR5 for 8 h, at which point 20 µM MB1^WDR5^ (competitor), or equal volume of buffer (control), were added. Fluorescence spectra were recorded 18 h later. T = 37 °C.

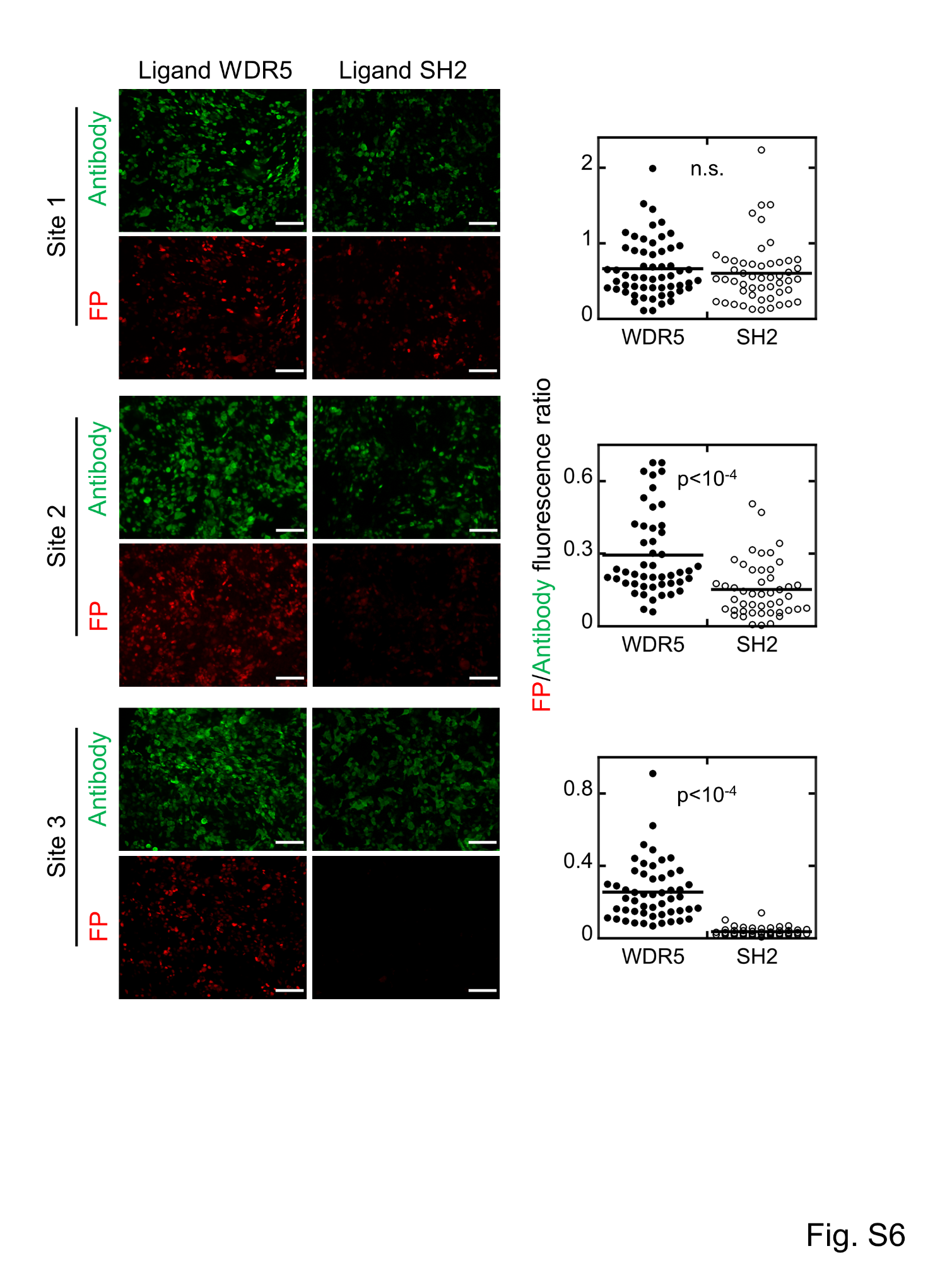

**Figure S6, related to Figure 4. Screening r-ATOM^WDR5^ in cells.** r-ATOM^WDR5^ sensors were composed of MB1^WDR5^ fused to TagRFP (non-circularly permuted) at site 1, site 2, or site 3. HEK 293T cells were co-transfected with one plasmid encoding r-ATOM^WDR5^ and a second plasmid expressing WDR5 or SH2. Cells were fixed, stained with anti-RFP antibody and imaged in green (antibody) and red (biosensor) channels. Scale bars are 100 µm. The ratio of intensities of the red and green channels for each ligand is shown at right. Each data point represents one cell. Significance was determined by a *t­*-test with unequal variance (n.s., not statistical). The results are representative of at least two biological repeats**.**

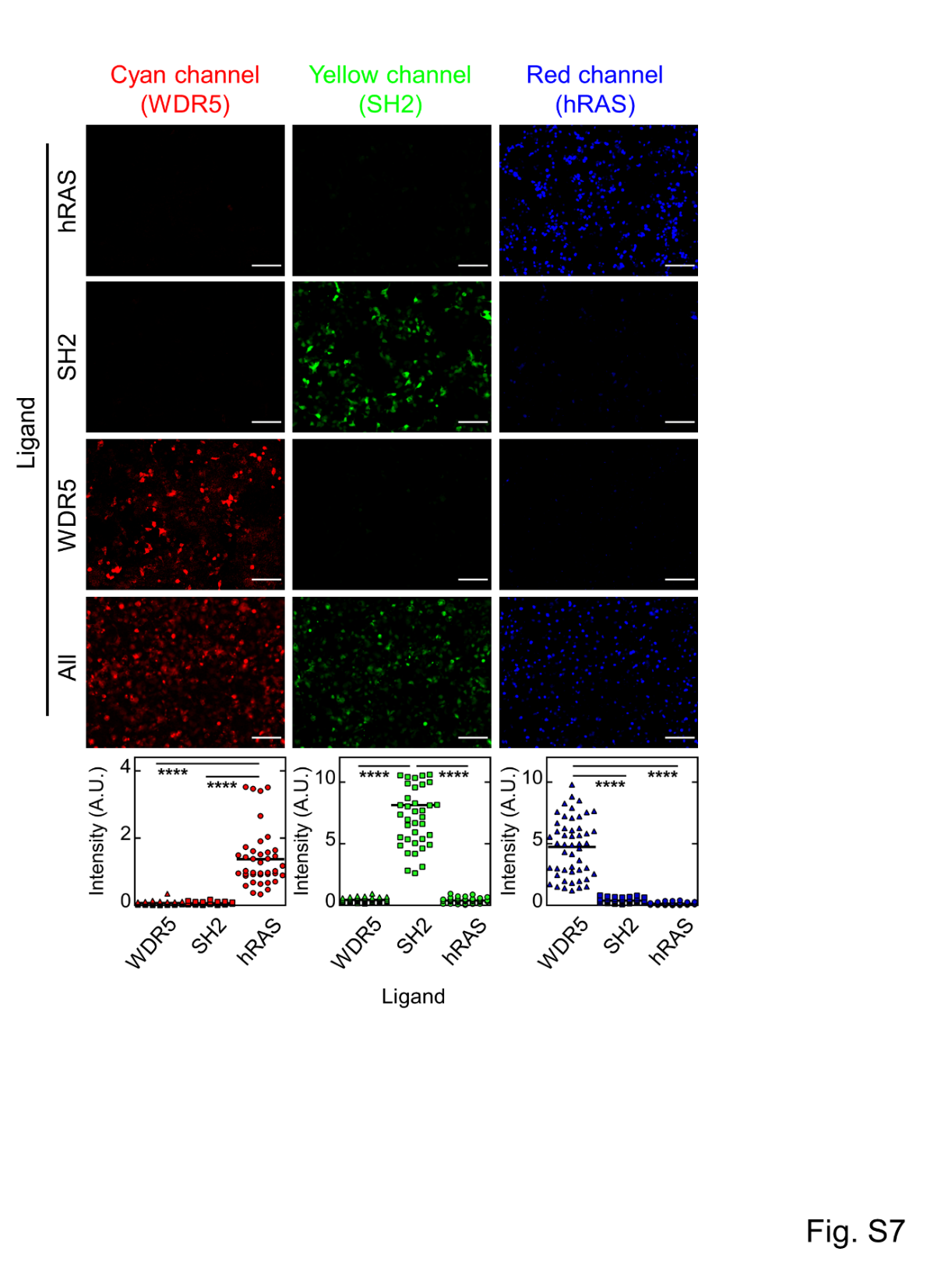

**Figure S7, related to Figure 4. r-ATOM^WDR5^, y-ATOM^SH2^, and c-ATOM^RAS^, co-expressed in the same cells, exhibit high turn-on values in the presence of their target ligands but not in the presence of noncognate ligands.** To match Figure 4, the cyan channel is represented in red, the yellow channel in green, and the red channel in blue. HEK 293T cells were co-transfected with an equimolar mixture of three biosensor plasmids and a fourth plasmid encoding hRAS (targeted to plasma membrane; top row), SH2 (expressed in cytoplasm; second row), or WDR5 (targeted to nucleus; third row). The fourth row shows the same experiment, but the fourth plasmid expressed all three ligands as described in the text. Scale bars are 100 µm. Turn-on ratios in each color channel were calculated by dividing the average intensity of cells expressing the biosensor and its cognate ligand by the average of the intensities of cells expressing the same biosensor and the two negative control ligands (bottom row). Turn-on values were 23-fold (r-ATOM^WDR5^), 19-fold (y-ATOM^SH2^), and 22-fold (c-ATOM^RAS^). Significance was determined by a *t­*-test with unequal variance: ****, p<10^-4^. The results are representative of three biological repeats**.**

y-ATOM^WDR5^

**MLPDNHYLSYQSVLSKDPNEGNSPVQKFKVPGSKSTATISGLKPGVDYTITVYAYQGGGRWHPYGYYSPISINYRTGGSGSGGASGGATGGSGGGSSVPTKLEVVAATPTSLLISWDAPAVTVVHYVITYGETGKRDHMVLLEFVTAAGITLGMDELYKGGGSGGMVSKGEELFTGVVPILVELDGDVNGHKFSVRGEGEGDATNGKLTLKLICTTGKLPVPWPTLVTTLGYGLACFSRYPDHMKQHDFFKSAMPEGYVQERTISFKDDGTYKTRAEVKFEGDTLVNRIELKGIDFKEDGNILGHKLEYNFNSHNVYITADKQKNGIKANFKIRHNVEDGSVQLADHYQQNTPIGDGPVLLEHHHHHHHH**

y-ATOM^SH2^

**MLPDNHYLSYQSVLSKDPNEKRDHMVLLEFVTAAGITLGMDELYKGGGSGGMVSKGEELFTGVVPILVELDGDVNGHKFSVRGEGEGDATNGKLTLKLICTTGKLPVPWPTLVTTLGYGLACFSRYPDHMKQHDFFKSAMPEGYVQERTISFKDDGTYKTRAEVKFEGDTLVNRIELKGIDFKEDGNILGHKLEYNFNSHNVYITADKSPVQEFTVPYSSSTATISGLSPGVDYTITVYAWGEDSAGYMFMYSPISINYRTGGSGSGGASGGATGGSGGGSSVPTKLEVVAATPTSLLISWDAPMSSSSVYYYRITYGETGGNQKNGIKANFKIRHNVEDGSVQLADHYQQNTPIGDGPVLLEHHHHHHHH**

RBP-y-ATOM^RAS^

**MKEGKTIGLVISTLNNPFFVTLKNGAEEKAKELGYKIIVEDSQNDSSKELSNVEDLIQQKVDVLLINPVDSDAVVTAIKEANSKNIPVITIDRSANGGDVVSHIASDNVKGGEMAAEFIAKALKGKGNVVELEGIPGASAARDRGKGFDEAIAKYPDIKIVAKQAADFDRSKGLSVMENILQAQPKIDAVFAQNDEMALGAIKAIEAANRQGIIVVGFDGTEDALKAIKEGKMAATIAQQPALMGSLGVEMADKYLKGEKIPNFIPAELKLITKENVQGGAASGGAAGGSSAARLQVDKAAAGLPDNHYLSYQSVLSKDPNEKRDHMVLLEFVTAAGITLGMDELYKGGGSGGMVSKGEELFTGVVPILVELDGDVNGHKFSVRGEGEGDATNGKLTLKLICTTGKLPVPWPTLVTTLGYGLACFSRYPDHMKQHDFFKSAMPEGYVQERTISFKDDGTYKTRAEVKFEGDTLVNRIELKGIDFKEDGNILGHKLEYNFNSHNVYITADKQKNGIKANFKIRHNVEDGVDYTITVYAWGWHGQVYYYMGSPISINYRTGGSGSGGASGGATGGSGGGSSVPTKLEVVAATPTSLLISWDAPAVTVDYYVITYGETGGNSPVQKFEVPGSKSTATISGLKPGSVQLADHYQQNTPIGDGPVLLEHHHHHHHH**

MBP-y-ATOM^WDR5^

**MKTEEGKLVIWINGDKGYNGLAEVGKKFEKDTGIKVTVEHPDKLEEKFPQVAATGDGPDIIFWAHDRFGGYAQSGLLAEITPDKAFQDKLYPFTWDAVRYNGKLIAYPIAVEALSLIYNKDLLPNPPKTWEEIPALDKELKAKGKSALMFNLQEPYFTWPLIAADGGYAFKYENGKYDIKDVGVDNAGAKAGLTFLVDLIKNKHMNADTDYSIAEAAFNKGETAMTINGPWAWSNIDTSKVNYGVTVLPTFKGQPSKPFVGVLSAGINAASPNKELAKEFLENYLLTDEGLEAVNKDKPLGAVALKSYEEELAKDPRIAATMENAQKGEIMPNIPQMSAFWYAVRTAVINAASGRQTVDEALKDAQTNSSSLEVLFQGPKVPGAATPVATMLPDNHYLSYQSVLSKDPNEGNSPVQKFKVPGSKSTATISGLKPGVDYTITVYAYQGGGRWHPYGYYSPISINYRTGGSGSGGASGGATGGSGGGSSVPTKLEVVAATPTSLLISWDAPAVTVVHYVITYGETGKRDHMVLLEFVTAAGITLGMDELYKGGGSGGMVSKGEELFTGVVPILVELDGDVNGHKFSVRGEGEGDATNGKLTLKLICTTGKLPVPWPTLVTTLGYGLACFSRYPDHMKQHDFFKSAMPEGYVQERTISFKDDGTYKTRAEVKFEGDTLVNRIELKGIDFKEDGNILGHKLEYNFNSHNVYITADKQKNGIKANFKIRHNVEDGSVQLADHYQQNTPIGDGPVL**

MBP-y-ATOM^SH2^

**MKTEEGKLVIWINGDKGYNGLAEVGKKFEKDTGIKVTVEHPDKLEEKFPQVAATGDGPDIIFWAHDRFGGYAQSGLLAEITPDKAFQDKLYPFTWDAVRYNGKLIAYPIAVEALSLIYNKDLLPNPPKTWEEIPALDKELKAKGKSALMFNLQEPYFTWPLIAADGGYAFKYENGKYDIKDVGVDNAGAKAGLTFLVDLIKNKHMNADTDYSIAEAAFNKGETAMTINGPWAWSNIDTSKVNYGVTVLPTFKGQPSKPFVGVLSAGINAASPNKELAKEFLENYLLTDEGLEAVNKDKPLGAVALKSYEEELAKDPRIAATMENAQKGEIMPNIPQMSAFWYAVRTAVINAASGRQTVDEALKDAQTNSSSLEVLFQGPKVPGAATPVTADMLPDNHYLSYQSVLSKDPNEKRDHMVLLEFVTAAGITLGMDELYKGGGSGGMVSKGEELFTGVVPILVELDGDVNGHKFSVRGEGEGDATNGKLTLKLICTTGKLPVPWPTLVTTLGYGLACFSRYPDHMKQHDFFKSAMPEGYVQERTISFKDDGTYKTRAEVKFEGDTLVNRIELKGIDFKEDGNILGHKLEYNFNSHNVYITADKSPVQEFTVPYSSSTATISGLSPGVDYTITVYAWGEDSAGYMFMYSPISINYRTGGSGSGGASGGATGGSGGGSSVPTKLEVVAATPTSLLISWDAPMSSSSVYYYRITYGETGGNQKNGIKANFKIRHNVEDGSVQLADHYQQNTPIGDGPVL**

Continued next page

MBP-y-ATOM^RAS^

**MKTEEGKLVIWINGDKGYNGLAEVGKKFEKDTGIKVTVEHPDKLEEKFPQVAATGDGPDIIFWAHDRFGGYAQSGLLAEITPDKAFQDKLYPFTWDAVRYNGKLIAYPIAVEALSLIYNKDLLPNPPKTWEEIPALDKELKAKGKSALMFNLQEPYFTWPLIAADGGYAFKYENGKYDIKDVGVDNAGAKAGLTFLVDLIKNKHMNADTDYSIAEAAFNKGETAMTINGPWAWSNIDTSKVNYGVTVLPTFKGQPSKPFVGVLSAGINAASPNKELAKEFLENYLLTDEGLEAVNKDKPLGAVALKSYEEELAKDPRIAATMENAQKGEIMPNIPQMSAFWYAVRTAVINAASGRQTVDEALKDAQTNSSSLEVLFQGPKVPGAATPVTADMLPDNHYLSYQSVLSKDPNEKRDHMVLLEFVTAAGITLGMDELYKGGGSGGMVSKGEELFTGVVPILVELDGDVNGHKFSVRGEGEGDATNGKLTLKLICTTGKLPVPWPTLVTTLGYGLACFSRYPDHMKQHDFFKSAMPEGYVQERTISFKDDGTYKTRAEVKFEGDTLVNRIELKGIDFKEDGNILGHKLEYNFNSHNVYITADKQKNGIKANFKIRHNVEDGVDYTITVYAWGWHGQVYYYMGSPISINYRTGGSGSGGASGGATGGSGGGSSVPTKLEVVAATPTSLLISWDAPAVTVDYYVITYGETGGNSPVQKFEVPGSKSTATISGLKPGSVQLADHYQQNTPIGDGPVL**

NES-MBP-c-ATOM^RAS^ (mutations from y-ATOM^RAS^ are highlighted)

**MNLVDLQKKLEELELDEQQGPVATMKTEEGKLVIWINGDKGYNGLAEVGKKFEKDTGIKVTVEHPDKLEEKFPQVAATGDGPDIIFWAHDRFGGYAQSGLLAEITPDKAFQDKLYPFTWDAVRYNGKLIAYPIAVEALSLIYNKDLLPNPPKTWEEIPALDKELKAKGKSALMFNLQEPYFTWPLIAADGGYAFKYENGKYDIKDVGVDNAGAKAGLTFLVDLIKNKHMNADTDYSIAEAAFNKGETAMTINGPWAWSNIDTSKVNYGVTVLPTFKGQPSKPFVGVLSAGINAASPNKELAKEFLENYLLTDEGLEAVNKDKPLGAVALKSYEEELAKDPRIAATMENAQKGEIMPNIPQMSAFWYAVRTAVINAASGRQTVDEALKDAQTNSSSLEVLFQGPKVPGAATPVTADMLPDNHYLSTQSVLSKDPNEKRDHMVLLEFVTAAGITLGMDELYKGGGSGGMVSKGEELFTGVVPILVELDGDVNGHKFSVRGEGEGDATNGKLTLKLICTTGKLPVPWPTLVTTLSWGVQCFARYPDHMKQHDFFKSAMPEGYVQERTISFKDDGTYKTRAEVKFEGDTLVNRIELKGIDFKEDGNILGHKLEYNYISDNVYITADKQKNGIKANFKIRHNVEDGVDYTITVYAWGWHGQVYYYMGSPISINYRTGGSGSGGASGGATGGSGGGSSVPTKLEVVAATPTSLLISWDAPAVTVDYYVITYGETGGNSPVQKFEVPGSKSTATISGLKPGSVQLADHYQQNTPIGDGPVL**

NLS-MBP-r-ATOM^WDR5^

**MPKKKRKVGGSGSMKTEEGKLVIWINGDKGYNGLAEVGKKFEKDTGIKVTVEHPDKLEEKFPQVAATGDGPDIIFWAHDRFGGYAQSGLLAEITPDKAFQDKLYPFTWDAVRYNGKLIAYPIAVEALSLIYNKDLLPNPPKTWEEIPALDKELKAKGKSALMFNLQEPYFTWPLIAADGGYAFKYENGKYDIKDVGVDNAGAKAGLTFLVDLIKNKHMNADTDYSIAEAAFNKGETAMTINGPWAWSNIDTSKVNYGVTVLPTFKGQPSKPFVGVLSAGINAASPNKELAKEFLENYLLTDEGLEAVNKDKPLGAVALKSYEEELAKDPRIAATMENAQKGEIMPNIPQMSAFWYAVRTAVINAASGRQTVDEALKDAQTNSSSLEVLFQGPKVPGAATPVATMVSKGEELIKENMHMKLYMEGTVNNHHFKCTSEGEGKPYEGTQTMRIKVVEGGPLPFAFDILATSFMYGSRTFINHTQGIPDFFKQSFPEGFTWERVTTYEDGGVLTATQDTSLQDGCLIYNVKIRGVNFPSNGPVMQKKTLGWEANTEMLYPADGGLEGRTDMALKLVGGGHLICNFKTTYRSKKPAKNLKMPGVYYVDHRLERIKEAD GNSPVQKFKVPGSKSTATISGLKPGVDYTITVYAYQGGGRWHPYGYYSPISINYRTGGSGSGGASGGATGGSGGGSSVPTKLEVVAATPTSLLISWDAPAVTVVLYVITYGETGKETYVEQHEVAVARYCDLPSKLGHKLN**

Prolactin secretion signal-y-ATOM^SH2^-KDEL

**MNIKGSPWKGSLLLLLVSNLLLCQSVAPVTADLPDNHYLSYQSVLSKDPNEKRDHMVLLEFVTAAGITLGMDELYKGGGSGGMVSKGEELFTGVVPILVELDGDVNGHKFSVRGEGEGDATNGKLTLKLICTTGKLPVPWPTLVTTLGYGLACFSRYPDHMKQHDFFKSAMPEGYVQERTISFKDDGTYKTRAEVKFEGDTLVNRIELKGIDFKEDGNILGHKLEYNFNSHNVYITADKSPVQEFTVPYSSSTATISGLSPGVDYTITVYAWGEDSAGYMFMYSPISINYRTGGSGSGGASGGATGGSGGGSSVPTKLEVVAATPTSLLISWDAPMSSSSVYYYRITYGETGGNQKNGIKANFKIRHNVEDGSVQLADHYQQNTPIGDGPVLGSGKDEL**

Cox4ss-y-ATOM^SH2^

**MLSLRQSIRFFKPATRTLCSSRYLLAPVTADLPDNHYLSYQSVLSKDPNEKRDHMVLLEFVTAAGITLGMDELYKGGGSGGMVSKGEELFTGVVPILVELDGDVNGHKFSVRGEGEGDATNGKLTLKLICTTGKLPVPWPTLVTTLGYGLACFSRYPDHMKQHDFFKSAMPEGYVQERTISFKDDGTYKTRAEVKFEGDTLVNRIELKGIDFKEDGNILGHKLEYNFNSHNVYITADKSPVQEFTVPYSSSTATISGLSPGVDYTITVYAWGEDSAGYMFMYSPISINYRTGGSGSGGASGGATGGSGGGSSVPTKLEVVAATPTSLLISWDAPMSSSSVYYYRITYGETGGNQKNGIKANFKIRHNVEDGSVQLADHYQQNTPIGDGPVL**

Continued next page

WDR5

**MHHHHHHSSGVDLGTENLYFQSNIGSGVPIGAVHGGHPGVVHPPQQPLPTAPSGPNSLQPNSVGQPGATTSSNSSASNKSSLSVKPNYTLKFTLAGHTKAVSAVKFSPNGEWLASSSADKLIKIWGAYDGKFEKTISGHKLGISDVAWSSDSRLLVSGSDDKTLKVWELSTGKSLKTLKGHSNYVFCCNFNPQSNLIVSGSFDESVRIWDVRTGKCLKTLPAHSDPVSAVHFNRDGSLIVSSSYDGLCRIWDTASGQCLKTLIDDDNPPVSFVKFSPNGKYILAATLDNTLKLWDYSKGKCLKTYTGHKNEKYCIFANFSVTGGKWIVSGSEDNMVYIWNLQSKEVVQKLQGHTDTVLCTACHPTENIIASAALENDKTIKLWKSDT**

SH2

**MPSNYITPVNSLEKHSWYHGPVSRNAAEYLLSSGINGSFLVRESESSPGQRSISLRYEGRVYHYRINTASDGKLYVSSESRFNTLAELVHHHSTVADGLITTLHYPAPKRNKPTVYGVSPNY**

hRAS

**MTEYKLVVVGAVGVGKSALTIQLIQNHFVDEYDPTIEDSYRKQVVIDGETCLLDILDTAGQEEYSAMRDQYMRTGEGFLCVFAINNTKSFEDIHQYREQIKRVKDSDDVPMVLVGNKCDLAARTVESRQAQDLARSYGIPYIETSAKTRQGVEDAFYTLVREIRQHLEHHHHHHHH**

hRAS-CAAX

**MTEYKLVVVGAVGVGKSALTIQLIQNHFVDEYDPTIEDSYRKQVVIDGETCLLDILDTAGQEEYSAMRDQYMRTGEGFLCVFAINNTKSFEDIHQYREQIKRVKDSDDVPMVLVGNKCDLAARTVESRQAQDLARSYGIPYIETSAKTRQGVEDAFYTLVREIRQHGGTSGSGGAGASGGSGTGGGASKLRKLNPPDESGPGCMSCKCVLS**

WDR5-P2A-SH2-P2A-hRAS-CAAX

**MVPIGAVHGGHPGVVHPPQQPLPTAPSGPNSLQPNSVGQPGATTSSNSSASNKSSLSVKPNYTLKFTLAGHTKAVSAVKFSPNGEWLASSSADKLIKIWGAYDGKFEKTISGHKLGISDVAWSSDSRLLVSGSDDKTLKVWELSTGKSLKTLKGHSNYVFCCNFNPQSNLIVSGSFDESVRIWDVRTGKCLKTLPAHSDPVSAVHFNRDGSLIVSSSYDGLCRIWDTASGQCLKTLIDDDNPPVSFVKFSPNGKYILAATLDNTLKLWDYSKGKCLKTYTGHKNEKYCIFANFSVTGGKWIVSGSEDNMVYIWNLQSKEVVQKLQGHTDTVLCTACHPTENIIASAALENDKTIKLWKSDTGLRSRAGSGATNFSLLKQAGDVEENPGPGSGPNSATMPSNYITPVNSLEKHSWYHGPVSRNAAEYLLSSGINGSFLVRESESSPGQRSISLRYEGRVYHYRINTASDGKLYVSSESRFNTLAELVHHHSTVADGLITTLHYPAPKRNKPTVYGVSPNYKLGSGATNFSLLKQAGDVEENPGPGSGGSMTEYKLVVVGAVGVGKSALTIQLIQNHFVDEYDPTIEDSYRKQVVIDGETCLLDILDTAGQEEYSAMRDQYMRTGEGFLCVFAINNTKSFEDIHQYREQIKRVKDSDDVPMVLVGNKCDLAARTVESRQAQDLARSYGIPYIETSAKTRQGVEDAFYTLVREIRQHGGTSGSGGAGASGGSGTGGGASKLRKLNPPDESGPGCMSCKCVLS**

Prolactin secretion signal-SH2-KDEL

**MNIKGSPWKGSLLLLLVSNLLLCQSVAPVTADPSNYITPVNSLEKHSWYHGPVSRNAAEYLLSSGINGSFLVRESESSPGQRSISLRYEGRVYHYRINTASDGKLYVSSESRFNTLAELVHHHSTVADGLITTLHYPAPKRNKPTVYGVSPNYGSGKDEL**

Cox4ss-SH2

**MLSLRQSIRFFKPATRTLCSSRYLLAPVTADPSNYITPVNSLEKHSWYHGPVSRNAAEYLLSSGINGSFLVRESESSPGQRSISLRYEGRVYHYRINTASDGKLYVSSESRFNTLAELVHHHSTVADGLITTLHYPAPKRNKPTVYGVSPNY**

**Figure S8. Amino acid sequences of ATOM biosensors and their ligands.** For ATOM sequences, MB domains are in blue, FP domains in green, circular permutation and other linker sequences in purple, and HisTag in grey. Other protein and peptide tags are colored or underlined as indicated.

| **Table S1: Quantification of cell images, related to Figures 1, 2, 4, and 5** | | | | |
| --- | --- | --- | --- | --- |
| **Figure** | **Description** | **# Cells** | **Average intensity** | **s.d.** |
| 1D | MB1^RAS^ FP site 1, (+) Ligand | 54 | 1.29 | 0.51 |
| 1D | MB1^RAS^ FP site 1, (-) Ligand | 51 | 1.91 | 0.53 |
| 1D | MB1^RAS^ FP site 2, (+) Ligand | 53 | 0.34 | 0.07 |
| 1D | MB1^RAS^ FP site 2, (-) Ligand | 53 | 0.52 | 0.07 |
| 1D | MB1^RAS^ FP site 3, (+) Ligand | 49 | 0.04 | 0.03 |
| 1D | MB1^RAS^ FP site 3, (-) Ligand | 54 | 0.05 | 0.01 |
| 1D | MB2^RAS^ FP site 1, (+) Ligand | 52 | 1.62 | 0.62 |
| 1D | MB2^RAS^ FP site 1, (-) Ligand | 53 | 1.13 | 0.32 |
| 1D | MB2^RAS^ FP site 2, (+) Ligand | 44 | 0.59 | 0.5 |
| 1D | MB2^RAS^ FP site 2, (-) Ligand | 43 | 0.09 | 0.05 |
| 1D | MB2^RAS^ FP site 3, (+) Ligand | 43 | 0.17 | 0.12 |
| 1D | MB2^RAS^ FP site 3, (-) Ligand | 36 | 0.09 | 0.03 |
| 1D | MB1^SH2^ FP site 1, (+) Ligand | 37 | 1.62 | 0.96 |
| 1D | MB1^SH2^ FP site 1, (-) Ligand | 56 | 0.69 | 0.26 |
| 1D | MB1^SH2^ FP site 2, (+) Ligand | 47 | 0.22 | 0.14 |
| 1D | MB1^SH2^ FP site 2, (-) Ligand | 59 | 0.12 | 0.05 |
| 1D | MB1^SH2^ FP site 3, (+) Ligand | 52 | 0.05 | 0.02 |
| 1D | MB1^SH2^ FP site 3, (-) Ligand | 45 | 0.04 | 0.01 |
| 1D | MB2^SH2^ FP site 1, (+) Ligand | 48 | 0.53 | 0.31 |
| 1D | MB2^SH2^ FP site 1, (-) Ligand | 48 | 0.39 | 0.13 |
| 1D | MB2^SH2^ FP site 2, (+) Ligand | 47 | 0.07 | 0.03 |
| 1D | MB2^SH2^ FP site 2, (-) Ligand | 59 | 0.08 | 0.02 |
| 1D | MB2^SH2^ FP site 3, (+) Ligand | 51 | 0.07 | 0.02 |
| 1D | MB2^SH2^ FP site 3, (-) Ligand | 50 | 0.05 | 0.01 |
| 1D | MB1^WDR5^ FP site 1, (+) Ligand | 53 | 2.79 | 0.62 |
| 1D | MB1^WDR5^ FP site 1, (-) Ligand | 53 | 2.86 | 0.85 |
| 1D | MB1^WDR5^ FP site 2, (+) Ligand | 41 | 0.24 | 0.09 |
| 1D | MB1^WDR5^ FP site 2, (-) Ligand | 44 | 0.12 | 0.04 |
| 1D | MB1^WDR5^ FP site 3, (+) Ligand | 53 | 0.35 | 0.19 |
| 1D | MB1^WDR5^ FP site 3, (-) Ligand | 41 | 0.07 | 0.02 |
| 2 | y-ATOM^WDR5^, WDR5 Ligand | 75 | 4.87 | 1.86 |
| 2 | y-ATOM^WDR5^, SH2 Ligand | 33 | 0.23 | 0.09 |
| 2 | y-ATOM^WDR5^, RAS Ligand | 35 | 0.26 | 0.09 |
| 2 | y-ATOM^SH2^, WDR5 Ligand | 66 | 0.25 | 0.08 |
| 2 | y-ATOM^SH2^, SH2 Ligand | 64 | 4.78 | 3.04 |
| 2 | y-ATOM^SH2^, RAS Ligand | 54 | 0.25 | 0.09 |
| 2 | y-ATOM^RAS^, WDR5 Ligand | 66 | 0.31 | 0.08 |
| 2 | y-ATOM^RAS^, SH2 Ligand | 53 | 0.29 | 0.08 |
| 2 | y-ATOM^RAS^, RAS Ligand | 62 | 4.99 | 2.48 |
| 4 | Cyan Channel (+) Ligand | 40 | 1.62 | 0.61 |
| 4 | Cyan Channel (-) Ligand | 41 | 0.07 | 0.03 |
| 4 | Yellow Channel (+) Ligand | 40 | 1.36 | 0.07 |
| 4 | Yellow Channel (-) Ligand | 41 | 0.1 | 0.04 |
| 4 | Red Channel (+) Ligand | 40 | 0.74 | 0.32 |
| 4 | Red Channel (-) Ligand | 41 | 0.12 | 0.05 |
| 5 | ER Sensor, Cytoplasmic Ligand | 63 | 0.08 | 0.04 |
| 5 | ER Sensor, ER Ligand | 52 | 0.77 | 0.31 |
| 5 | ER Sensor, Mitochondrial Ligand | 55 | 0.06 | 0.02 |
| 5 | Mito Sensor, Cytoplasmic Ligand | 58 | 0.05 | 0.02 |
| 5 | Mito Sensor, ER Ligand | 54 | 0.04 | 0.02 |
| 5 | Mito Sensor, Mitochondrial Ligand | 51 | 0.88 | 0.62 |

| **Table S2: Quantification of cell images, related to Supporting Figures S4, S6, and S7** | | | | |
| --- | --- | --- | --- | --- |
| **Figure** | **Description** | **# Cells** | **Average intensity** | **s.d.** |
| S4 | Antibody stain: y-ATOM^WDR5^ with WDR5 Ligand | 61 | 647 | 164 |
| S4 | Antibody stain: y-ATOM^WDR5^ with SH2 Ligand | 48 | 647 | 220 |
| S4 | Antibody stain: y-ATOM^WDR5^ with RAS Ligand | 50 | 691 | 286 |
| S4 | Biosensor Fluorescence: y-ATOM^WDR5^ with WDR5 Ligand | 61 | 116 | 67 |
| S4 | Biosensor Fluorescence: y-ATOM^WDR5^ with SH2 Ligand | 48 | 6.9 | 2.7 |
| S4 | Biosensor Fluorescence: y-ATOM^WDR5^ with RAS Ligand | 50 | 6.5 | 2.9 |
| S4 | Antibody stain: y-ATOM^SH2^ with WDR5 Ligand | 49 | 954 | 368 |
| S4 | Antibody stain: y-ATOM^SH2^ with SH2 Ligand | 49 | 875 | 271 |
| S4 | Antibody stain: y-ATOM^SH2^ with RAS Ligand | 51 | 823 | 223 |
| S4 | Biosensor Fluorescence: y-ATOM^SH2^ with WDR5 Ligand | 49 | 55 | 28 |
| S4 | Biosensor Fluorescence: y-ATOM^SH2^ with SH2 Ligand | 49 | 1056 | 494 |
| S4 | Biosensor Fluorescence: y-ATOM^SH2^ with RAS Ligand | 51 | 64 | 32 |
| S4 | Antibody stain: y-ATOM^RAS^ with WDR5 Ligand | 50 | 446 | 169 |
| S4 | Antibody stain: y-ATOM^RAS^ with SH2 Ligand | 47 | 423 | 194 |
| S4 | Antibody stain: y-ATOM^RAS^ with RAS Ligand | 47 | 622 | 273 |
| S4 | Biosensor Fluorescence: y-ATOM^RAS^ with WDR5 Ligand | 50 | 9.5 | 7.4 |
| S4 | Biosensor Fluorescence: y-ATOM^RAS^ with SH2 Ligand | 47 | 12.1 | 7.8 |
| S4 | Biosensor Fluorescence: y-ATOM^RAS^ with RAS Ligand | 47 | 180 | 93 |
| S6 | r-ATOM^WDR5^ Site 1 with WDR5 Ligand | 57 | 0.66 | 0.38 |
| S6 | r-ATOM^WDR5^ Site 1 with SH2 Ligand | 51 | 0.60 | 0.41 |
| S6 | r-ATOM^WDR5^ Site 2 with WDR5 Ligand | 48 | 0.29 | 0.17 |
| S6 | r-ATOM^WDR5^ Site 2 with SH2 Ligand | 47 | 0.17 | 0.11 |
| S6 | r-ATOM^WDR5^ Site 3 with WDR5 Ligand | 52 | 0.25 | 0.16 |
| S6 | r-ATOM^WDR5^ Site 3 with SH2 Ligand | 49 | 0.03 | 0.02 |
| S7 | Cyan Channel with WDR5 Ligand | 51 | 0.06 | 0.05 |
| S7 | Cyan Channel with SH2 Ligand | 43 | 0.05 | 0.03 |
| S7 | Cyan Channel with RAS Ligand | 40 | 1.37 | 0.85 |
| S7 | Yellow Channel with WDR5 Ligand | 50 | 0.42 | 0.14 |
| S7 | Yellow Channel with SH2 Ligand | 43 | 8.12 | 3.51 |
| S7 | Yellow Channel with RAS Ligand | 38 | 0.39 | 0.21 |
| S7 | Red Channel with WDR5 Ligand | 51 | 4.65 | 2.4 |
| S7 | Red Channel with SH2 Ligand | 43 | 0.38 | 0.19 |
| S7 | Red Channel with RAS Ligand | 40 | 0.13 | 0.11 |
